# TransferGWAS: GWAS of images using deep transfer learning

**DOI:** 10.1101/2021.10.22.465430

**Authors:** Matthias Kirchler, Stefan Konigorski, Matthias Norden, Christian Meltendorf, Marius Kloft, Claudia Schurmann, Christoph Lippert

## Abstract

**Motivation:** Medical images can provide rich information about diseases and their biology. However, investigating their association with genetic variation requires non-standard methods. We propose *transferGWAS*, a novel approach to perform genome-wide association studies directly on full medical images. First, we learn semantically meaningful representations of the images based on a transfer learning task, during which a deep neural network is trained on independent but similar data. Then, we perform genetic association tests with these representations.

**Results:** We validate the type I error rates and power of *transferGWAS* in simulation studies of synthetic images. Then we apply *transferGWAS* in a genome-wide association study of retinal fundus images from the UK Biobank. This first-of-a-kind GWAS of full imaging data yielded 60 genomic regions associated with retinal fundus images, of which 7 are novel candidate loci for eye-related traits and diseases.

**Supplementary information:** Our method is implemented in Python and available at https://github.com/mkirchler/transferGWAS/

## 1 Introduction

The emergence of large-scale imaging-genetics datasets and deep learning provides us with the potential for powerful genome-wide association studies (GWAS) using image phenotypes. Current approaches to GWAS of imaging data have been largely focused on extracting fixed quantities of interest from the images, such as organ volumes, distances, and sizes. Such quantities have been obtained manually (e.g., segmenting organ volumes using graphics software), by automated software tools [16], or deep learning models trained to predict segmentation masks or other biomarkers [21, 15, 35, 41, 3, 17]. However, such methods are limited in the scope of questions they can address and prohibit the detection of associations between unspecified variation of the studied organ and the genetic variation. Further challenges are that (i) deep learning methods require large amounts of labeled data, which can be very costly to produce, (ii) the extracted sets of biomarkers are generally study- and disease-specific [8, 36], and (iii) deep-learning-based feature extraction often leads to noisier estimates of the trait under investigation compared to hand-labeled and quality-controlled labels [18]. Overall, it is unclear how to aggregate the complex geometric and topological structures present in images into a form beyond traditional biomarkers and useful in a GWAS setting. While imaging GWAS based on specific extracted image features is valuable and important for scientific discovery, here we present a complementary approach called *transferGWAS*. Instead of extracting pre-specified phenotypes from images, *transferGWAS* treats the images themselves as phenotypes. This can be achieved with a version of transfer learning, which is well-established in deep learning. By training a deep neural network on an independent dataset and extracting transformations of the original images at hidden layers of the network, we obtain low-dimensional, semantically meaningful representations [43]. These representations are continuous variables and can be analyzed using association tests for quantitative traits. A related line of work has sought to find general associations between histology images and genetic variants and gene expression using a deep-learning based version of canonical correlation analysis (CCA) [2, 19]. However, the respective methods have not yet been adapted for a GWAS setting. We evaluate type I error rate and power of *transferGWAS* in Monte Carlo simulation studies, where, conditional on genetic effects, we generate images using generative adversarial networks (GAN). We apply *transferGWAS* to retinal fundus images from the UK Biobank [7] and detail ways to understand and interpret the discovered associations.

## 2 System and Methods

*transferGWAS* is a multi-step method for using imaging data as phenotypes in GWAS. At its core, study images are condensed into lower-dimensional feature embeddings using a deep convolutional neural network (CNN) trained on an independent “transfer” dataset. CNNs can learn complex non-linear and layered continuous representations of the input data. Early layers usually extract simple information similar to edge detectors, while deeper layers learn more complex features [43, 44]. In each layer of the network, the representations become more sophisticated but also more specific to the learning task [44]. Empirical evidence supports that deep neural networks learn useful representations of their input data that is transferable even to widely diverging tasks [43]. We show that the learned representations are to some degree task-agnostic, meaning that even though the CNN has to be trained on a specific task (such as predicting disease states from images), the representations capture considerably more information than what is necessary to perform the specific transfer task. In the context of two-sample testing, a recent study has shown both theoretically and empirically that a similar transfer-hypothesis testing approach yields valid p-values and exhibits high statistical power [24].

*transferGWAS* consists of three main steps (Fig. 1). First, a CNN is trained on an auxiliary dataset, called the “transfer dataset.” The transfer dataset can depict the same image modality as the study images in the next step but can also consist of non-medical images. At this stage, no genetic data is used. In the second step, the dataset of interest is analyzed, which must be independent of the transfer dataset regarding the observations. Using the CNN, the images (“study images”) available for the participants in the GWAS cohort are condensed into continuous low-dimensional phenotype representations. For this, the images are fed into the network trained in the first step, after which the results are extracted at an intermediate, hidden layer of the network (instead of at the output layer) and then further reduced to ten dimensions via principal component analysis. Finally, genetic association testing is performed on the condensed phenotypes using linear mixed models. All steps can be adapted to fit the investigator’s interests, and especially customization of the first two steps allows exploring different aspects of the data and indirect specification of prior knowledge.

**Figure 1:**
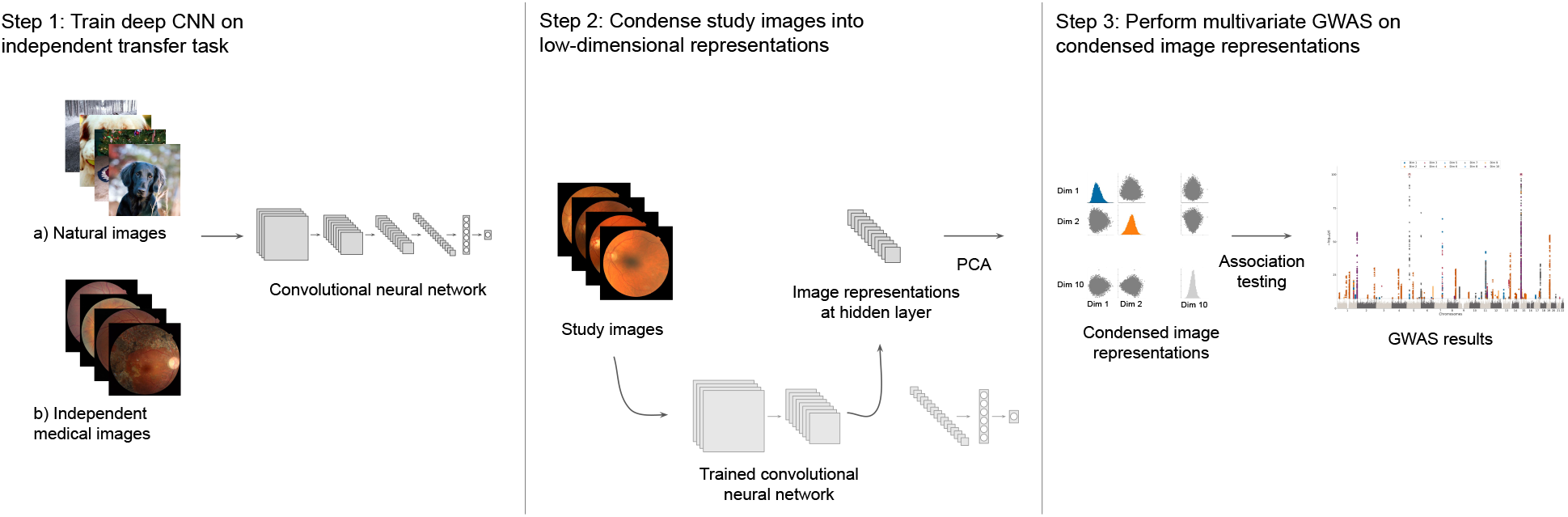
Overview of the *transferGWAS* approach which consists of three steps. First, a convolutional neural network is trained on an independent transfer task to prime the network. Second, *transferGWAS* uses the trained network and a principal component analysis to condense study images into low-dimensional embeddings. Last, a linear mixed model association analysis is performed on the image representations.

Two main mechanisms set *transferGWAS* apart from classical GWAS approaches. First, *transferGWAS* allows the implicit modeling of hard-to-model known phenotypes, as neural networks can efficiently encode complex patterns with minimal information loss between image and hidden representation [13]. Second, neural networks can also encode previously unknown phenotypical patterns. The pre-training stage primes a neural network either to task-specific or to general features – potentially uncovering novel phenotypic variations that domain experts are unaware of.

### 2.1 Pre-training

We evaluate two different pre-training strategies. In both cases, we use the same convolutional neural network architecture, a ResNet50 [20]. First, we use a ResNet50 trained on the ImageNet dataset of everyday objects (provided by the PyTorch library [33]). Second, we train another ResNet50 with randomly initialized weights on the EyePACS dataset of retinal fundus images [10]. We perform this training in a multi-task manner to predict different stages of diabetic retinopathy (DR) and reconstruct the original image from the bottleneck layer simultaneously. Both tasks share a common CNN body, but while for the DR prediction task the head of the network is a simple linear layer, for the image reconstruction task the head is a convolutional decoder network as in an autoencoder (Supp. Fig. 6). This allows the network to be primed to disease-related aspects while capturing as much information about the original image as possible. Although the network is explicitly trained to detect DR, the hidden layer embeddings are also expected to capture very different characteristics. This is partly because the network needs to distinguish DR from other eye diseases efficiently, and hence implicitly learns versatile eye attributes. For training the DR model, we use the Adam optimizer [23] with learning rate set to 10^*−* 3^, batch size 25, and the mean squared error loss function for both reconstruction and DR prediction (weighed equally). We train the network for 100 epochs, using the OneCycle learning rate schedule (using the weights from the epoch with the lowest validation loss). To avoid overfitting, images are randomly rotated, color-jittered, and flipped during each training step (“data augmentation”).

### 2.2 Feature Condensation

The ResNet50 architecture is organized into one simple convolutional layer at the beginning and four subsequent multi-layer blocks, usually also called layers. We extract image representations from the fourth (i.e., last) and second of the larger layer blocks for the ImageNet and EyePACS-trained networks, respectively. In both cases, we use the representation at the end of the layer block directly before applying the last batch normalization and rectified linear unit (ReLU) activation. We perform eight deterministic image transformations (rotation by 0°, 90°, 180°, and 270°, each both with and without a horizontal mirroring), feed all images individually into the network, and stack the outputs into a single feature vector. This is a version of test-time augmentation (TTA) shown to improve the robustness of deep learning models on prediction tasks [25]. For each individual, we stack the corresponding feature vectors of both left and right eye. Since we extract features at convolutional layers, the corresponding embeddings contain spatial information, which we average out (“global average pooling”). At this stage, the dimensionality of the feature representations is n_channels *×* 8 *×* 2, where n_channels is 512 for the second layer block and 2048 for the last layer block of a ResNet50 architecture. We further reduce the dimensionality via a principal component analysis to the first ten PCs, which accounted for 52.5% of the explained variance on the ImageNet-trained network. For the EyePACS-trained network, the first ten PCs accounted for about 98.6% of the explained variance of the full embedding phenotype; see Supp. Fig. 7a and Sec. 2.4.

### 2.3 Genetic Association Testing

We perform genetic association tests in a linear mixed model (LMM) framework. There are different possible approaches to handle multivariate phenotypic data in an LMM framework, such as multi-variate LMMs [45, 26] or other approaches [1]. In this paper, we perform a univariate association test for each of the ten feature dimensions from Sec. 2.2 and then compute the minimum p-value of all ten dimensions, Bonferroni-corrected by a factor of 10 to adjust for multiple testing of the ten feature dimensions. This procedure controls the family-wise error rate and is conservative since it does not account for the relationships between different feature dimensions. We use the infinitesimal model with leave-one-chromosome-out scheme in the BOLT-LMM software package [30, 29, 27]. To adjust for confounding, we include age, sex, indicator variables for genotyping microarray and assessment center, and the first ten genetic PCs as provided by the UK Biobank as covariates. Most feature dimensions were approximately normally distributed, but a small number of dimensions were skewed in one direction, violating the Gaussianity assumption of BOLT-LMM. To counteract this effect, we apply a rank-based inverse normal transformation (INT) [4] for all feature dimensions before fitting the LMM. We first compute residuals from fitting only the fixed-effect covariates (age, sex, genotyping array, assessment center, and first ten genetic PCs) in a linear regression model, adjust those residuals with the INT and finally use BOLT-LMM on those adjusted residuals while again correcting for the covariates, as proposed by [38]. Notably, the association results were virtually identical with and without INT adjustment on the first ten feature dimensions both for ImageNet- and EyePACS-pre-training. However, for a small number of higher-order feature dimensions, the INT suppressed many likely false-positive association results (Sec. 2.4).

We perform the *transferGWAS* on the 16,255,985 SNPs from the imputed genotype data provided by the UKB (Supp. Sec. 1.3). To estimate the genetic relationship matrix, we only use a subset of the directly genotyped variants, which accounted for 557,891 SNPs (BOLT-LMM recommends 500k genotypes) after filtering for MAF ≥ 0.1%, Hardy-Weinberg equilibrium (significance level: 0.001), pairwise LD pruning (*R*^2^=0.8) and maximum missingness of 10% per SNP and individual. For LD Score regression to calibrate BOLT-LMM test statistics, we used the European-ancestry scores supplied by the library. For interpretation, we group associated variants in regions, which were defined as 500kb genomic segments carrying at least one associated SNP with p-value*<* 10^*−* 9^.

### 2.4 Hyperparameter Selection in *transferGWAS*

For the ImageNet-trained network, we select the penultimate layer (after the last convolution operation, before batch normalization and ReLU activation), as is common in many transfer settings. Since ImageNet contains 1,000 different classes, the resulting image representation is sufficiently diverse and general to yield useful results. For the CNN trained on the EyePACS dataset, the training task is very narrow. To find more than only diabetic-retinopathy related traits, we choose the end of the second layer (of the four convolutional layer-blocks in the ResNet50 architecture).

We select other hyperparameters based on a parameter sweep in a simulation study, designed analogously to the validation study for empirical type I error rate and power (Sec. 3). The number of principal components had relatively little influence on the power in the simulation study (as long as it was at least 5 PCs), and we fix it to ten. We validate this behavior on the real datasets (Supp. Fig. 7b). Without correcting skewed distributions via INT (Sec. 2.3), we found (in data not shown here) that a single feature dimension (#47) in EyePACS pre-training had a large number of likely false-positive SNP hits. Those were mostly one-or two-SNP loci with MAF *<* 0.5% and relatively low imputation quality, spread over 19 different chromosomes. These clusters of likely false-positive hits disappear when correcting the feature dimensions with INT while giving similar association results for other feature dimensions. The remaining hyperparameters we select to maximize empirical power in the simulation hyperparameter sweep. They were: image size (set to 448); whether or not to use test-time augmentation (set to use test-time augmentation); and whether to pool the spatial output at the convolutional layers via maximum or mean pooling (set to mean). We also considered using the hyperparameter sweep to select the optimal layer. However, the best layer for simulated images might very well not coincide with the best layer in the realistic scenarios of interest. This is because image features generated by the StyleGAN2 might operate on different spatial scales and can not be expected to capture significant pathologic patterns.

### 2.5 Interpretation of Black-box Results

Association results returned by *transferGWAS* cannot directly be traced back to specific phenotypic abstractions. To overcome this drawback, we apply several explanatory approaches to interpret the association results. First, we visually inspect the individual feature dimensions by contrasting images that strongly activate the respective dimensions in either direction. Next, we cross-correlate each individual feature dimension with all phenotypic traits collected in the UKBiobank in a phenome-wide association study (PheWAS, Suppl. Sec. 2.7) and compare association results to previous findings in the GWAS catalog [6]. Finally, we can also explore individual association results by training a CNN to predict the genotype of a specific SNP and then contrasting high- and low-activating images for this network. Namely, we train a ResNet18 architecture to predict the genotype from the left retinal fundus image. Images are resized to 448 *×* 448 pixels and randomly rotated by up to 20° during training. Assuming an additive genetic model, we code labels continuously as 0, 1, and 2. We use the Pearson correlation coefficient between predictions and true genotypes in each mini-batch as loss function. Using ImageNet pre-training, we further train the network for 10 epochs (with early stopping) with Adam optimizer and OneCycle learning rate scheduler, learning rate of 0.0001, and a batch-size of 64. We visualize the images in the validation dataset that most activate in negative or positive direction after training.

## 3 Method Validation in Simulation Study

To investigate the empirical type I error rate and power of *transferGWAS*, we perform a novel Monte Carlo simulation study. We generate synthetic images conditioned on 512-dimensional latent “code vectors.” These code vectors, in turn, are simulated from a small set of “causal” SNPs. We choose simulated causal SNPs uniformly at random from the first half of each chromosome (to estimate power) and count SNPs on the second half of each chromosome as null SNPs to evaluate type I error rate. To counteract proximal contamination [28] of the null SNPs, we additionally leave a gap of 2 Mb before and after the center of each chromosome. The effect size of each causal SNP is drawn from a standard normal distribution and then normalized such that all causal SNPs together account for the specified variation in every single dimension of the code vectors. We fix the number of causal SNPs to 1,250 across all chromosomes (out of 275,142 SNPs in total on the first halves of all chromosomes), vary the explained variation of latent embeddings between 0.5 and 0.7, and also vary sample size between 6,000 and 46,731. We replicate each setting 10 times with different random seeds for a total of 80 genome-wide simulation runs. See Supp. Tab. 2 for an overview of the scenarios. For the generation of synthetic images, we use the state-of-the-art StyleGAN2 [22], a deep learning architecture for synthesizing high-resolution images from low-dimensional latent codes. We train StyleGAN2 on a subset of healthy retinal fundus images from the EyePACS dataset, an independent dataset of retinal fundus images used to detect DR [10], using normally-distributed latent variables conditional on genetic variants as input. Then we perform association tests between the genetic variants and the synthetic images using *transferGWAS* with a ResNet50 architecture trained on ImageNet.

### 3.1 Results of Simulation Study

Supp. Fig. 1 shows a random selection of images generated by the StyleGAN2. The images generally appear realistic. There is a small number of synthetic retinal images with no or two optic disks. The GAN also exhibits some amount of “mode collapse”: several images look very similar, most images are of the left eye, and a central artifact can be seen in most images. In the explored simulation settings under the null hypothesis of no association between genetic variants and image phenotypes, the type I error rate was properly controlled: only two null SNPs reached genome-wide significance at *α* = 5 · 10^*−* 8^ (i.e., a type I error rate of 2.5%). Supp. Fig. 2a shows a QQ-plot of null SNPs for one exemplary simulation run. For analysis of the empirical power, we selected 1,250 causal SNPs, which accounted for 50% explained variance of the latent code. 12% of the 1,250 causal SNPs could be recovered with a sample size of 46,731 individuals. With 70% explained variance, the number of discovered causal SNPs increased to 24% (Supp. Fig. 2b and Supp. Tab. 2).

## 4 Results

We present results from applying *transferGWAS* on retinal fundus images of 46,731 participants of European ancestry from the UK Biobank, see Supp. Tab. 3 for descriptive statistics and Supp. Sec. 1.3 for details on the dataset and quality control. To characterize the information captured by the different feature dimensions, an experienced ophthalmologist visually explored those retinal fundus images which “activated” the extracted feature dimensions the most in positive and negative directions.

### 4.1 Feature Characterization

For the visual characterization of the features, Fig 2 and Supp. Fig. 3 and 4 show those images that activate each of the 10 feature dimensions the most in positive and negative direction, based on the ImageNet pre-training and the EyePACS pre-training, respectively. The most apparent characteristics captured by the feature dimensions based on ImageNet pre-training are brightness and hue, and structural attributes such as local smoothness. Besides this, some clinically relevant disease states are visible. For example, in dimension 6, the images activating in negative direction indicate severe myopic fundus, while in dimension 9, the images activating in positive direction show typical signs of an atrophy of the retinal pigment epithelial layer. The visual observation of the images activating the features based on the EyePACS pre-training show considerably more disease-related characteristics. The positive-activating images of dimension 1 show signs of pigment deposits of the retinal pigment epithelial layer, which can be caused by different eye pathologies such as retinitis pigmentosa. In the positive-activating images for dimensions 3 and 4 and negative-activating images for dimension 8, again, typical signs of a myopic fundus are visible, while the negative-activating images in dimension 4 also exhibit emphasized choroidal vessels. The positive-activating images for dimension 7 show strong indications of peripheral retinal drusen. For dimension 10, opacities in the vitreous humor as seen in pathologies like asteroid hyalosis can be discerned in both eyes. The remaining dimensions (2, 5, 6, and 9) do not show clear pathological patterns. Dimensions 2 and 6 seem to capture low-quality and noise signals, such as opacities and overexposure. Note that the images were recorded without dilation, which may explain the large number of exposure and shadow artifacts.

**Figure 2:**
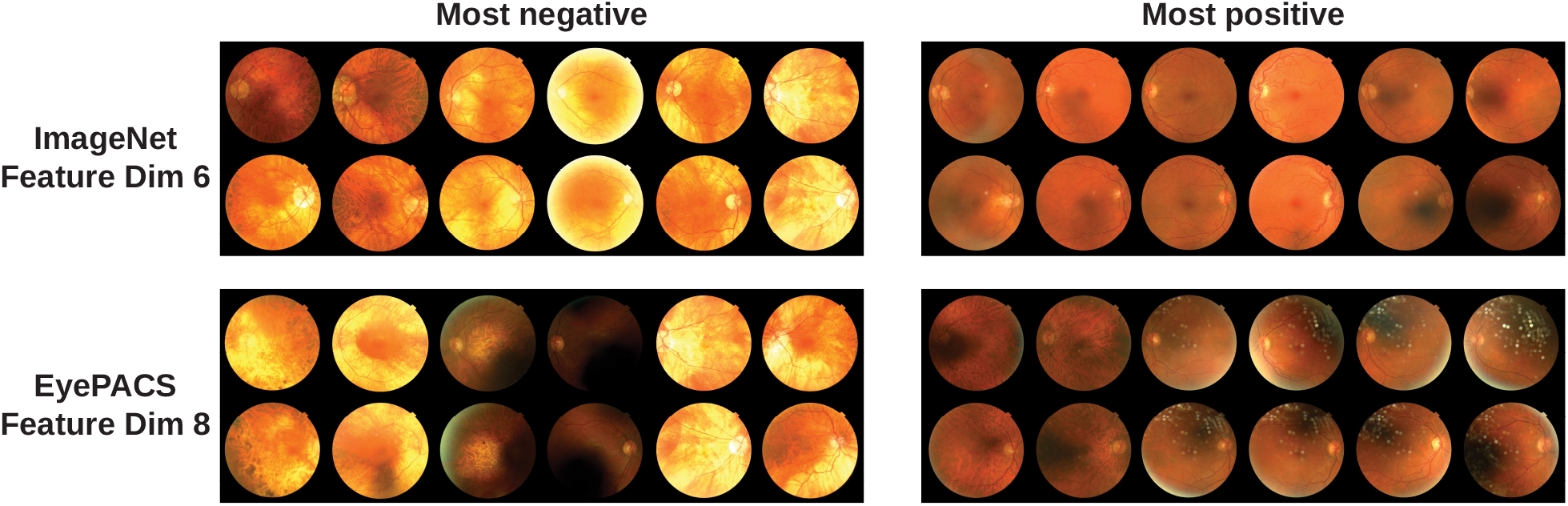
Illustration of information in the retinal fundus images captured by the feature dimensions. Shown are example images activating 2 selected feature dimensions: dimension 6 from ImageNet pre-training and dimension 8 from EyePACS pre-training. The remaining feature dimensions are shown in Supp. Fig. 3 and 4. Retinal fundus images are displayed for the left eye (upper row) and the right eye (lower row); left column (“most negative”) shows the six individuals activating the corresponding feature dimension the most in negative direction; right column (“most positive”) shows the six individuals activating the corresponding feature dimension the most in positive direction. Reproduced by kind permission of UK Biobank ©.

The PheWAS results are shown in Supp. File 1. For ImageNet pre-training and EyePACS pre-training, there were 205 and 266 significantly associated traits (Bonferroni-corrected). Each feature dimension was associated with between 19 and 124 traits (ImageNet pre-training) and between 1 and 139 traits (EyePACS pre-training). 179 of these traits were associated in both pre-trainings. Shared traits were related to ophthalmic information, containing specific eye diseases, history of eye surgery, results of different eye measurement procedures, and reasons for wearing glasses. In addition, associations with pigmentation traits, cardiometabolic traits, and lifestyle factors but also with technical parameters, blood cell counts, or lung function measurements can be observed. Of the 26 associated traits unique to ImageNet pre-training, two were eye disease/eye procedure traits, and three were other disease/medication-related traits. On the other hand, the 87 traits uniquely identified after the EyePACS pre-training included 21 eye disease/eye procedure traits and 12 other disease/medication-related traits. For a more specific comparison and accounting for various overlaps of the phenotypic traits, we focused on the reasons for wearing glasses and doctor-diagnosed eye problems and diseases as recorded by touchscreen questionnaires (Supp. Tab. 1). All 14 traits were associated with at least one feature dimension based on the EyePACS pre-training, and 9 traits were associated with at least one feature dimension based on the ImageNet pre-training.

### 4.2 GWAS

There was no evidence of unaccounted stratification in the *transferGWAS* (genomic control parameter *λ*_GC_ ranged between 0.999-1.087). We identified 48 and 31 regions associated with at least one feature dimension in the ImageNet and EyePACs pre-training, respectively. Using PLINK’s clumping method, we identified 95 and 63 independent loci located in the 48 and 31 regions for ImageNet and EyePACS, respectively. They account for 117 signals that can be mapped to 60 genomic regions. Manhattan plots are shown in Fig. 3a and 3b, QQ-plots in Supp. Fig. 5; for the full list of associated loci, see Supp. Files 2 and 3. We identified 19 regions that were significantly associated in both analyses based on the ImageNet and EyePACs pre-training. Furthermore, 23 regions were significant in one analysis and showed sub-threshold significance (p-value *<* 10^*−* 4^) in the other analysis. Additionally, we found 16 and 2 regions exclusively associated with at least one feature dimension in the ImageNet and EyePACs pre-training, respectively.

**Figure 3:**
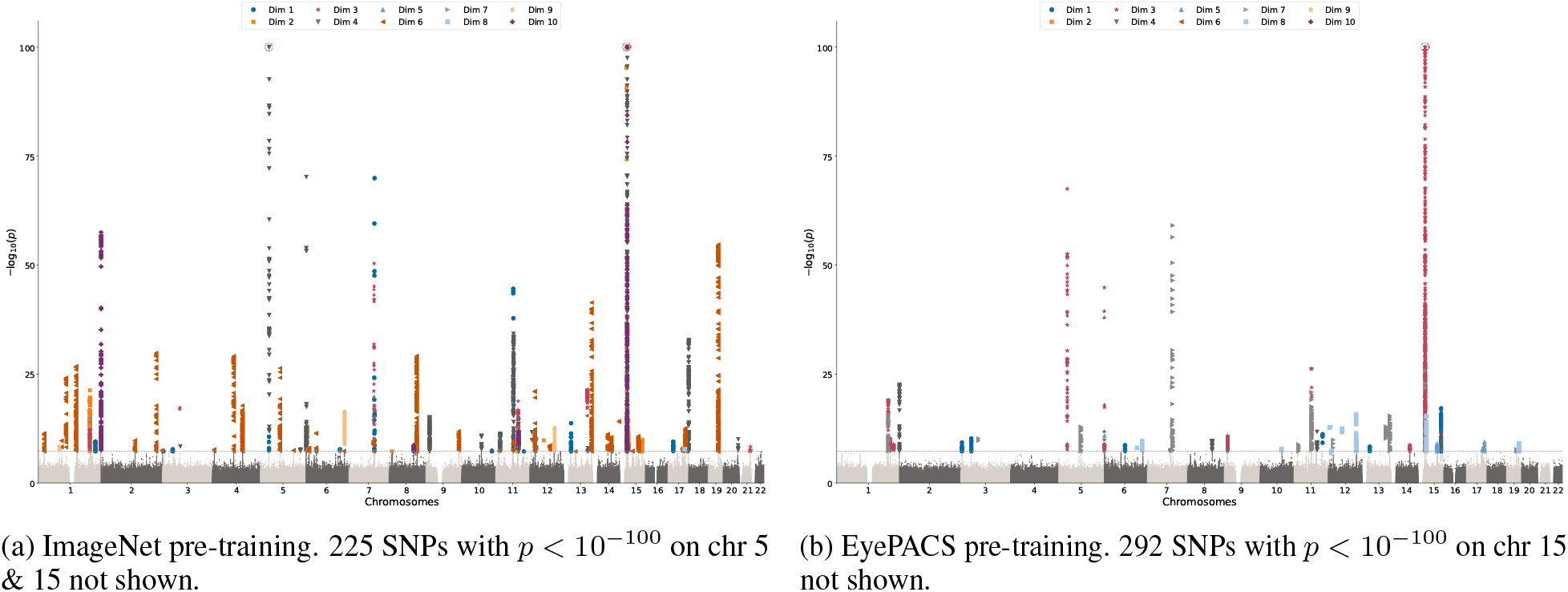
Manhattan plots for *transferGWAS* (*n* = 46, 731 individuals, 16,255,985 SNPs). Shown are -log_10_-transformed p-values across chromosomes 1 through 22. For each SNP, the minimum p-value over the ten feature dimensions multiplied by 10 (Bonferroni correction) is plotted. For SNPs exceeding the significance threshold (5 · 10^*−* 8^) the highest activating dimension is indicated by the corresponding marker. P-values below 10^*−* 100^ are truncated and indicated with a red circle for visualization purposes.

Inspecting the known associations for the 60 regions in the GWAS catalog, we found associations with eye diseases and related traits (including myopia, age-related macular degeneration, diabetic retinopathy, glaucoma, cataract, macular thickness, refractive error, intraocular pressure, corneal and optic nerve measurements) for 36/60 regions. For example, 8 of the regions had previously been associated with myopia. Of them, 7 were associated with ImageNet feature dimensions 2, 4, 6 or 10, and 6 of those were associated with EyePACS feature dimensions 4, 7 or 8, thereby supporting our PheWAS findings where the ImageNet and EyePACS feature dimensions 6 and 8, respectively, showed the strongest correlation with myopia. To further evaluate the myopia top hits, we performed a separate GWAS for myopia in UK Biobank using the same set of samples as for the *transferGWAS*. Out of 70 known myopia loci [39], we could replicate 46% in the myopia GWAS and 32% and 23% in the ImageNet and EyePACS *transferGWAS*, respectively, using Bonferroni correction (p-value *<* 0.05*/*70). Of the 12 genome-wide significant regions from our myopia GWAS, we found that 75% of them showed significant associations in both *transferGWAS* after Bonferroni correction (p-value *<* 0.05*/*12) and 17-25% reached genome-wide significance.

Inspecting the known associations for the 60 regions in the GWAS catalog further, 30 regions have previously been associated with anthropometric traits (e.g., body weight, height, waist-hip ratio, or waist circumference), 29 with cardiovascular and metabolic traits (e.g., blood pressure, lipid levels, hypertension, and coronary artery disease), 9 with diabetes and related traits, 7 with smoking status, and 5 with Alzheimer’s disease or dementia. Beyond that, a large proportion of the regions (*N* = 17*/*60) has previously been reported to be associated with eye and hair color and skin pigmentation, for example, the OCA2-HERC2 region. Overall, 17 regions did not have previously reported associations with eye-related traits and diseases or pigmentation traits, and 7 regions did not have any previously reported associations with traits listed above.

The GOLGA8M-APBA2 region (significantly associated in EyePACS and ImageNet analysis) is located close to the OCA2-HERC2 region and has previously been shown to be associated with hair color. Using stepwise conditional analyses, we identified 3 independent signals within that region (Supp. Tab. 4). Interestingly, the APBA2 gene encodes for a neuronal adapter protein expressed in the brain that interacts with the Alzheimer’s disease amyloid precursor protein (APP). Low levels of amyloid-beta proteins are biomarkers of Alzheimer’s disease and can be detected in the eye [42]. Alternatively, the detection of loci associated with Alzheimer’s disease may be due to artifacts because of failure to comply with instructions during image capture. Another interesting locus that has not previously been reported is the FOXF2 locus (EyePacs, lead SNP rs9405472, intronic variant, MAF = 0.4279, p-value = 2.0 · 10^*−* 11^). FOXF2 is mainly expressed in the lung, colon, and esophagus and has been implicated in anterior segment dysgenesis [32]. Further genes in that region include additional genes from the forkhead family of transcription factors, including FOXC1, which plays a role in ocular development and has been implicated in glaucoma [37].

To better understand the found associations, we trained separate CNNs to predict the respective genetic variant from the fundus images (Fig 4). First, we investigated the GNB3 locus with lead SNP rs5442. rs5442 is a missense variant (MAF = 0.070) in GNB3 whose minor allele A has been shown to be associated with a 30% increased risk of high myopia [5]. Images activating in direction of the risk allele A show myopic fundus, which was not present in G-allele direction. Second, we inspected the ACTN4 locus with lead SNP rs10415219 (MAF = 0.486). In the A-allele activating images, retinal venular tortuosity is clearly visible, a potential marker for risk factors for cardiovascular diseases [9]. rs10415219 is in LD with rs1808382 (*R*^2^ = 1), whose major allele has previously been reported to be associated with increased retinal venular tortuosity [40] which is in line with our findings. For other selected SNPs, the visual inspection was not always conclusive.

**Figure 4:**
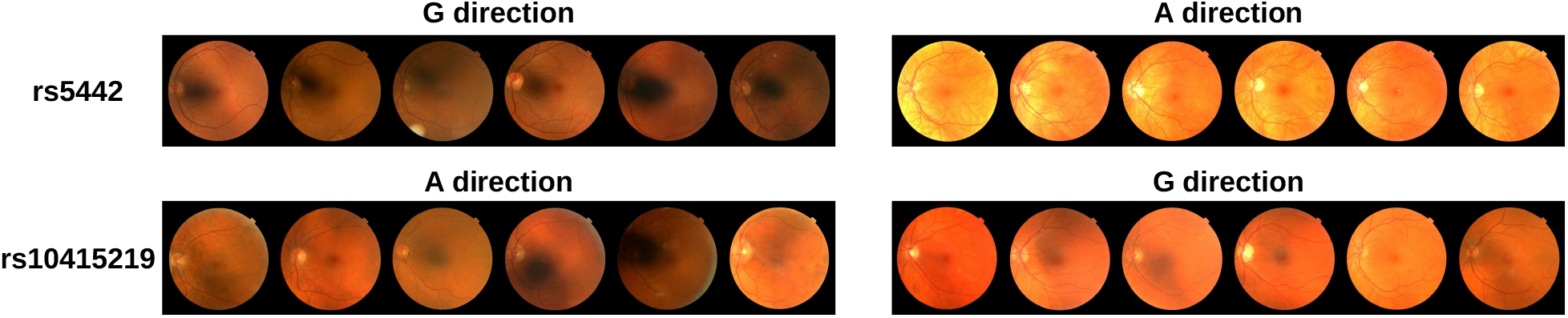
Visualization of two exemplary lead SNPs. Only left eyes are shown and were used for training. rs5442 shows strong signs of myopic fundus in the A direction, while rs10415219 shows retinal venular tortuosity in A direction. Images are plotted according to the predictions of a specifically trained CNN, but true genotypes may differ. Reproduced by kind permission of UK Biobank ©.

To assess if *transferGWAS* can yield results that would not be uncovered by a baseline GWAS performed on standard phenotypes, we selected three sets of standard phenotypes collected in the UKB: (1) directly eye-related traits, such as eye diseases; (2) all of the before and all traits indicated as “physical measures,” such as blood pressure; and (3) all of the before and all traits indicated as “health and medical history.” After quality-control and binarization of categorical traits, 92, 170, and 335 traits remained, respectively. For each of these 335 traits, we performed a GWAS with BOLT-LMM with the same settings and on the same subset of individuals as for our *transferGWAS*. Due to compute constraints, we only compare results from non-imputed genetic data in both *transferGWAS* and the baseline GWAS (yielding fewer loci than in the main analysis). The ImageNet and EyePACS *transferGWAS* yielded 74 and 26 total independent loci, while the baseline GWAS yielded 128, 221, and 223 loci, respectively, for the three sets of phenotypes. However, we found that the loci between *transferGWAS* and the baseline GWAS only intersect in 2-3 loci each (Supp. Tab. 5 and Supp. File 4), i.e., almost all of the loci uncovered by *transferGWAS* would not have been found by the baseline GWAS on standard traits, and vice versa. This indicates that at the same sample size, transferGWAS does indeed uncover genetic associations that standard GWAS approaches do not.

For example, rs2472299 was identified by the ImageNet *transferGWAS* (*p* = 3 · 10^*−* 10^) and was previously identified (via proxy SNP rs936226, *R*^2^ = 0.81) for systolic & diastolic blood pressure in a set of 187k individuals (140k of which from the UKB; *p* = 3 · 10^*−* 16^ and 10^*−* 18^ for SBP/DBP) [14]. In the baseline GWAS, however, rs2472299 only achieved p-values of down to 7 · 10^*−* 4^ for SBP & DBP (which was also the smallest p-value of all 335 traits for this SNP).

To further investigate how much signal the images carry that is not contained in the standard traits, we performed conditional transferGWAS, where we adjust for a large number of phenotypical traits in addition to the confounding factors. In particular, we adjust for several eye-related problems, cardiovascular traits, and pigmentation traits. We find that *transferGWAS* adjusted for the additional traits still yields 65 (ImageNet) and 38 (EyePACS) independent loci (on the imputed genetic data), a reduction of only 31.6% - 39.7% compared to the unadjusted *transferGWAS*. Note that we would expect the adjustment for many variables to cause a drop in power even when adjusting for pure noise variables. Among the remaining association results, many regions have previously been associated to the respective adjustment variables (eye-related problems, cardiovascular traits, and pigmentation). This further corroborates that at identical sample size, *transferGWAS* can capture signal with respect to these traits that is not encoded in the variables themselves.

## 5 Discussion

In the application of *transferGWAS* to the UK Biobank, we identified 60 genetic regions associated with retina scans, including 7 novel candidate loci. The remaining loci contain signals for eye-related traits and diseases, anthropometric, cardiovascular and metabolic traits, smoking status, and Alzheimer’s disease. One intriguing example of the power of *transferGWAS* is the identification of loci on chromosome 15 in the OCA2-HERC2 locus, for which there is evidence of their implication in oculocutaneous albinism [11, 12]. Oculocutaneous albinism exhibits general symptoms which provides a straightforward explanation of why *transferGWAS* can identify it in retina scans by linking pigmentation and eye diseases. Other novel loci proposed by *transferGWAS* have also been linked to disease-relevant processes and are interesting candidates for follow-up studies, such as the APBA2 locus on chromosome 15. The signals within that locus might represent novel independent loci. One possible explanation is the interaction of the translated protein of APBA2 with the amyloid precursor protein APP, which is involved in Alzheimer’s disease and related to eye diseases [42]. Importantly, we found that *transferGWAS* yielded almost only hits that could not be found in baseline GWAS on standard phenotypes at the same sample size. Even after adjusting the association analysis for many standard phenotypes, *transferGWAS* still retains a large fraction of its power. Both observations indicate that deep-learning-based image embeddings capture at least partially orthogonal information that is not present in the standard phenotypes and may potentially help close the missing heritability gap for complex traits [31].

We also demonstrated how combining PheWAS results, re-training of CNN classifiers for discovered SNPs, and subsequent visual inspection can reveal clinically relevant features such as myopic fundus and retinal venular tortuosity. Our results also confirm previous findings that a wide variety of physical signals and functions can be detected from retinal images [34].

Comparing the two different pre-training approaches, the specialized pre-training on EyePACS yielded feature dimensions that capture more pronounced eye-specific and eye-disease specific information compared to ImageNet pre-training. In the GWAS, however, ImageNet pre-training resulted in considerably more association results. This apparent contradiction is probably due to the more general training task for ImageNet pre-training. Although EyePACS pre-training captures more information about eye diseases, ImageNet pre-training identifies more non-specific information related to traits such as pigmentation and cardiometabolics. In the UK Biobank disease-specific phenotypes are overrepresented compared to non-pathological traits; e.g., “eye color” is not recorded at all, while most diseases are redundantly recorded in multiple fields. The choice of pre-training method in practice depends on the researcher’s goals – e.g., whether the analysis should be targeted to a specific disease group or more general traits are also of interest – and the imaging modality and available transfer datasets. Both a simulation study and a pilot study on a small subset can help gauge the comparative advantages without invalidating association results on the rest of the data.

We also presented a novel way to validate imaging-based GWAS using synthetically generated images. In Monte Carlo simulation studies, we generated images conditional on genetic variants to investigate the empirical type I error rate and power of *transferGWAS*. This evaluation helped to validate *transferGWAS* and indicated that hypothesis testing within *transferGWAS* can be made with valid type I error control.

### 5.1 Conclusions

We presented *transferGWAS*, a novel approach to utilize full images in GWAS, yielding interesting insights that might be easily overlooked in GWAS of pre-selected biomarkers extracted from images. We showed how *transferGWAS* can incorporate prior expert knowledge and illustrated its use for the analysis of retinal fundus images. Besides this use case, *transferGWAS* can be extended to any imaging modality. It can be used on 2D image (e.g., histological tissue images), 3D volumetric image (e.g., MR images of the brain), or CINE data (e.g., MR videos of the cardiac cycle). Even more generally, our method can be adapted to work on other complex biomarkers, such as graphs (e.g., brain connectomes) and time-series (e.g., ECG or EEG).

## Supporting information

Supplementary File 1 - PheWAS Results

Supplementary File 2 - GWAS Results

Supplementary File 3 - Regional Plots

Supplementary File 4 - Baseline Comparison

## Acknowledgements

We thank the participants of the UK Biobank study. This research has been conducted using the UK Biobank resource. This work was supported by the German Ministry of Research and Education (BMBF) [project numbers 01|S21069A, 01IS18051A, 031B0770E, and 01IS21010C]; the German Research Foundation (DFG) [project numbers KL 2698/2-1 and KL 2698/5-1]; and by the Carl-Zeiss Foundation to MKloft.

