## Supplementary Text, Tables, and Figures for "TransferGWAS: GWAS of images using deep transfer learning"

**This file includes:**

Dataset Specifics
Details of UK Biobank Analysis
Additional Simulation Results
Legends for Supplementary Files 1-4
Supplementary Figures 1-7
Supplementary Tables 1-6

#### 1 Dataset Specifics

In the analyses, we used three different datasets: The ImageNet and EyePACS image datasets were used in pre-training steps; a subset of healthy individuals of EyePACS were used to create synthetic images in the simulation study; and the UK Biobank dataset was used for the application, imaging GWAS, and the genetic data in the simulation study.

##### 1.1 ImageNet

ImageNet is a large-scale database of annotated images, sorted into one of many hierarchically structured classes. In this work we use the common "Imagenet Large Scale Visual Recognition Challenge" (ILSVRC2012, [Russakovsky et al., 2015]) subset, which consists of 1.2 million images in 1,000 classes. Among these classes are a wide variety of animals, plants, and everyday objects and devices.

##### 1.2 EyePACS

During the pre-training step for the follow-up analysis we use the EyePACS dataset [Cuadros and Bresnick, 2009]. The dataset was published in a Kaggle challenge, with the aim to build prediction models for diabetic retinopathy. The images were provided by EyePACS, which is a platform for retinopathy screening. The dataset consists of 88,702 retinal fundus images (44,351 individuals, one image for each eye) exhibiting different stages of diabetic retinopathy as well as healthy controls. Images came from different models and types of cameras and were recorded among several distinct sites. Images were center-cropped before any transformations were applied and then resized to $448 \times 448$ pixels.

##### 1.3 UK Biobank

The UK Biobank is a large-scale cohort study with rich genotypic and phenotypic profiles of about 500,000 participants of the general population from Great Britain [Bycroft et al., 2018]. The dataset contains genetic, imaging, and rich phenotype data including cardiometabolic and eye-related traits. Age, hair color, skin color, tanning, smoking behaviour, and information regarding eye-related conditions were self-reported via a touchscreen interface in the visit of the study participants in the assessment center. BMI was computed from the height and weight measured in the assessment center as kg/m2. Systolic and diastolic blood pressure were measured twice in the assessment center automatically in sitting position, and were averaged for the analysis. All data analyzed here was from the first visit to the assessment center (see Supplementary Table 3). In addition, information about main and secondary disease diagnoses were available from hospital admission data in the form of ICD10 codes. We used the UK Biobank as main dataset focusing on those individuals that self-identified as "White British" and had a similar genetic ancestry based on a genotype principal component analysis, in order to minimize the possibility of confounding due to population stratification. We found no outliers in pairwise scatter-plots of the first five genetic principal components.

Genetic data was available for 488,000 participants. Imputed genotype data was filtered according to minor allele frequency (MAF $\geq 0.1\%$), Hardy-Weinberg equilibrium (significance level: $10^{-12}$), imputation quality (information score $\geq 0.4$) as well as SNP and sample missingness of maximal $10\%$, which left 16,255,985 SNPs for analysis.

Retinal fundus imaging was performed using a Topcon 3D OCT-1000 Mark II system, with a field angle of 45 degrees, and without dilation. Data was recorded both between 2006-2010 and between 2012-2013 [Chua et al., 2019]. Using a dataset of retinal fundus images with annotated image quality [Fu et al., 2019], we trained a CNN to filter out low-quality images. In addition to this filtering step we visually inspected the remaining images and removed any images that were still dominated by strong noise features. After additionally filtering for missing covariates and genetic data, 46,731 white British participants remained for analysis.

### 2 Details of UK Biobank Analysis

#### 2.1 Identification of Independent GWAS Loci

The identified genome-wide significant variants from step 3 described above were mapped to independent signals using a $\pm 250$ kilo base pair (kb) window and an LD cutoff of $R^2 > 0.1$ using the clumping algorithm implemented in PLINK (p-value $< 10^{-9}$ for lead SNPs, p-value $< 5 \cdot 10^{-8}$ for associated SNPs).

#### 2.2 Lookup of Known GWAS Loci

For the GWAS catalog [Buniello et al., 2019] lookups we used the LDtrait API [Machiela and Chanock, 2015]. For each locus we looked up all known LD-linked associations for all SNPs in the locus with a window size of 250kb and $R^2 = 0.1$ (we did not use any p-value thresholds). The GWAS catalog was accessed on October 8, 2020.

#### 2.3 Comparison with Myopia GWAS

We performed an additional GWAS with myopia as an outcome, on 46,704 of the 46,731 individuals in UK Biobank for which self-reported “Reason for wearing glasses: myopia” (UKB Code 6147) was available, with 30% myopia cases (small differences to numbers in Supplementary Table 1 are due to the way that PHESANT defines cases and missing values). We used BOLT-LMM with the same covariates as before, since it reports reliable inference with case fraction of at least $10\%$ for variants with MAF $> 0.1\%$. For comparison, we use the list of 72 myopia loci identified in [Tedja et al., 2018]. We selected proxies for two SNPs that were not in the imputed genetic data provided by UK Biobank ($R^2 = 1$ for both). One other SNP was monoallelic in the white British population, and one further SNP was not listed in the 1000G reference panel and was left out as well, leaving 70 lead SNPs for which we investigated replication based on the myopia GWAS, as well as *transferGWAS* with ImageNet and EyePACS pre-training.

#### 2.4 Conditional Analysis

In the conditional analysis, we investigated whether variants within a region constituted independent signals. For this, we included the SNP with the lowest remaining p-value in the region as a covariate and re-ran the LMM association study, iterating this procedure until no significant SNPs remained within the region. Individuals with low imputation certainty in the SNP used for adjustment were removed from the respective analyses.

#### 2.5 Comparison against Baseline GWAS on Standard Traits

We selected three sets of phenotypes from the UK Biobank for the baseline GWAS. The first set contains all traits whose path includes the word “eye.” This includes a large number of physical measures and self-reported conditions regarding eyesight and diseases. In the second set, in addition to the eye traits, we included all traits that were in the “Physical Measures” subcategory in UKB, including many anthropometric and cardiovascular traits. In the last set, we added all self-reported conditions from the “Health and Medical History” category, which contains broad summaries of the individuals’ health state.

For quality control, we only included traits that have been reported in the Neale Lab phenome-wide GWAS results on UKB (2nd round, http://www.nealelab.is/uk-biobank/). We only included continuous, integer, and categorical (single and multiple) variables, and categorical variables were one-hot-recoded into binary variables. For multi-categorical variables, negative values (e.g. "do not know", "prefer not to answer") were dropped as in the Neale Lab GWAS.

Due to computational constraints, GWAS were only performed on the microarray data instead of the imputed data, and we only compared against the results on microarray data for *transferGWAS*.

For each of the three sets of phenotypes we computed the minimum p-value after Bonferroni-adjustment for the number of traits. Identification of independent loci were performed as in the main analysis. We mapped significant loci between baseline GWAS and *transferGWAS* against each other by checking if the lead SNPs were in a ±250kb window of each other.

##### 2.6 *transferGWAS* with Additional Phenotypic Adjustments

The *transferGWAS* adjusted for additional phenotypes was performed with identical settings to the main analysis on the imputed genetic data. We selected the following variables for adjustment (in addition to the confounding variables sex, assessment center, genotyping batch, age, and genetic PCs 1-10):

*Pigmentation traits:* Skin color; Hair Color; Tanning.

*Eye traits:* Ever had eye surgery; current eye infection; wears glasses; eye problems; reasons for wearing glasses; spherical power; cylindrical power.

*Cardiovascular & anthropometric traits:* diabetes; smoking status; vascular heart problems; diastolic & systolic blood pressure; BMI.

##### 2.7 Details on the Phenome-wide Association Study

For the PheWAS, we use the software tool PHESANT [Millard et al., 2017] to perform automated phenome-wide association tests between all traits available in UK Biobank and the 10 extracted feature dimensions (after INT correction, Sec. 2.3). Association test results were available for 11,644 tested traits, including personal characteristics, lifestyle factors, body and organ measurements, and ICD10 codes from hospital visits. The association tests are based on linear, ordered logistic, multinomial, or logistic regression models depending on the scale of the outcome variable, with covariates age, sex, and assessment center. We use a Bonferroni correction to account for the $11,644 \cdot 10$ tests. For the characterization of the features and to compare the two different pre-trainings, we highlighted and manually explored unique and shared traits that showed significant associations. Due to the many different data categories (e.g., questionnaires, verbal interviews, physical measures, biological samples) some of the traits are overlapping and provide similar information. Therefore, we focus on two specific eye-related questionnaires in the dataset and repeat the association tests. We explicitly select the questionnaires on "Eye problems/disorders" and "Reasons for wearing glasses" to evaluate and compare the plausibility of the information captured by the different features.

#### 3 Additional Simulation Results

In a preliminary second simulation experiment, we generated another set of images together with a number of standard traits to compare the power of *transferGWAS* against a baseline GWAS on the standard traits. In this setting, we use causal and independent SNPs from the microarray SNP data as in the main simulation study. We draw 1,250 causal SNPs at random collected in a matrix $G \in \{0,1,2\}^{N\times 1250}$, and use a weight matrix $W \in \mathbb{R}^{512\times 1250}$ for a simulated "physiology" trait $P = \lambda WG + (1-\lambda)\epsilon_P$ with standard normally distributed noise $\epsilon \sim \mathcal{N}_{N,512}(0, I)$. The weights in $W$ are drawn from a standard normal distribution, but set to 0 with a probability of 0.95 and $\lambda \in (0, 1)$ is selected such that $WG$ accounts for 50% of the variance of $P$. $P$ represents the general physiology of the individual. Images are directly generated using the same StyleGAN2 as in the main simulation study from this physiology trait $P$. For comparison, from the physiology $P$, we also compute a set of 10 standard traits: $T = \alpha VP + (1-\alpha)\epsilon_T$, where $V \in \mathbb{R}^{512\times t}$ is again a weight matrix with standard normal entries and sparsity varying between 0.5, 0.8, and 0.95, and $t$ is the number of simulated traits ($t \in \{1, 10\}$). $\alpha$ was selected such that $VP$ accounted for an explained variance of $T$ between $20\%$ and $50\%$.

We performed *transferGWAS* on the simulated images and baseline GWAS with the same settings on the simulated traits for three random seeds. In this setting, our *transferGWAS* achieved an empirical power of $15.5\%$(±0.40% SEM). Results for the simulated phenotypes are collected in Supplementary Table 6.

**Data Availability**

The individual-level genotype, covariates, and image data are available by application from the UK Biobank (https://www.ukbiobank.ac.uk/register-apply/). The images used for training are publicly available from http://image-net.org/index (ImageNet dataset) and https://www.kaggle.com/c/diabetic-retinopathy-detection (EyePACS diabetic retinopathy dataset).

**Code Availability**

Code and trained neural networks underlying all analyses, including the simulation study, are available at https://github.com/mkirchler/transferGWAS. For the GWAS we used the BOLT-LMM library [Loh et al., 2015, 2018], and for the phenome-wide association scans we used the PHESANT software library [Millard et al., 2017], both of which are publicly available.

#### Supplementary Files

**Supplementary File 1. Detailed summary of the PheWAS results.** Summary of PheWAS results for all associated traits of the UK Biobank and the 10 feature dimensions after ImageNet and EyePACS pre-training.

**Supplementary File 2. Detailed summary of the GWAS results.** GWAS results based on ImageNet and EyePACS pre-training, with filtering SNPs based on imputation quality (INFO) score > 0.8 and minor allele frequency (MAF) > 1

**Supplementary File 3. Regional plots for genomic regions.** For each independent signal, a regional plot is shown. Multi-signal regions are indicated by the same color. The full results can also be viewed on LocusZoom: ImageNet pre-training: `https://my.locuszoom.org/gwas/711563/?token=772bd3eac15a4cbfbd74d52abb551649` EyePACS pre-training: `https://my.locuszoom.org/gwas/429077/?token=59a345fd50d143d1a97f4631d1088adb`

**Supplementary File 4: Comparison against standard baseline GWAS.** Significant GWAS loci for each of the three trait groups (eye-specific, eye-specific & physical measures, all), as well as the *transferGWAS* results. All results only on the non-imputed data.

#### Supplementary Figures

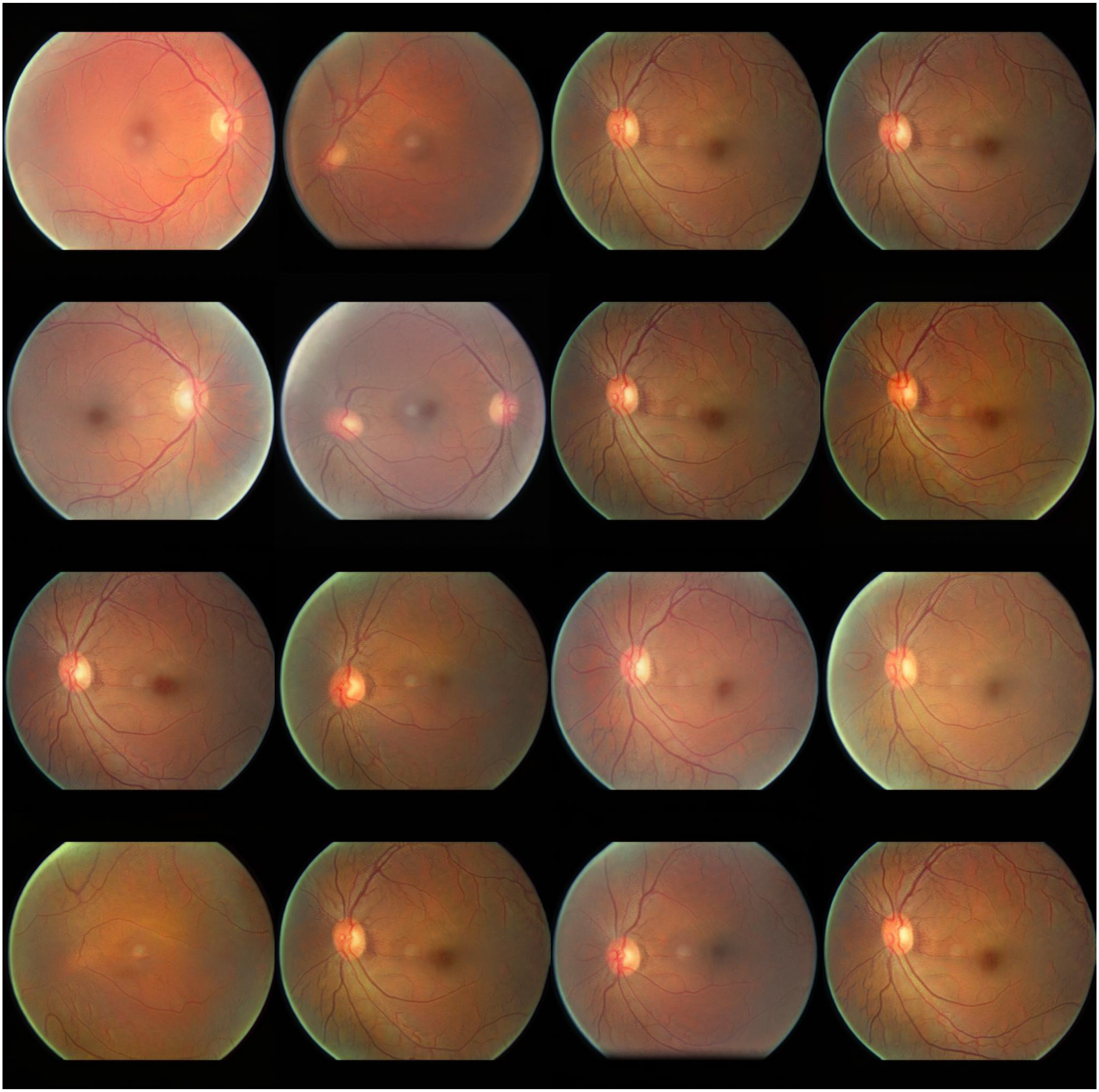

**Supplementary Figure 1. Illustration of synthetic images generated in the simulation study.**
Randomly selected synthetic images generated in the simulation study using the StyleGAN2. These images largely correspond to real fundus images, although some unnatural artifacts are visible (such as vasal abnormalities and alterations of the optic disc and macula vessels).

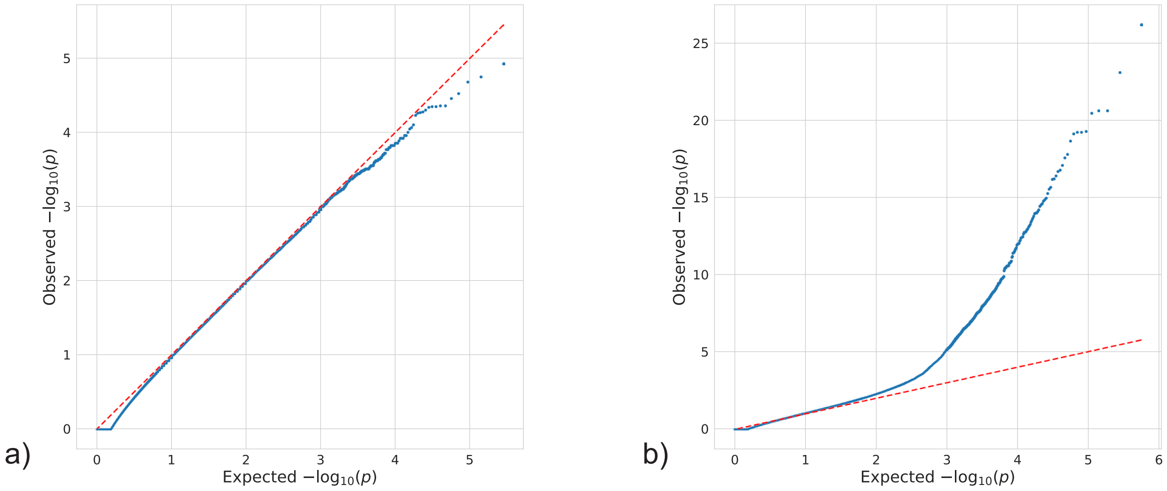


**Supplementary Figure 2. Distribution of the p-values from the simulation GWAS.** Shown are quantile-quantile plots of observed versus expected uniform p-values for one simulation run (n=46,731, explained variance of latent code=0.5, 1,250 causal SNPs), on the -log10 scale. P-values are Bonferroni-corrected minimum p-values over ten feature dimensions. a) shows only null SNPs (i.e. p-values of the SNPs on the second halves of all chromosomes); b) shows the p-values of all SNPs. The high number of SNPs with p-value of 1 is caused by the Bonferroni correction for 10 tests (i.e. multiplication by 10) and clipping p-values to the interval [0, 1].

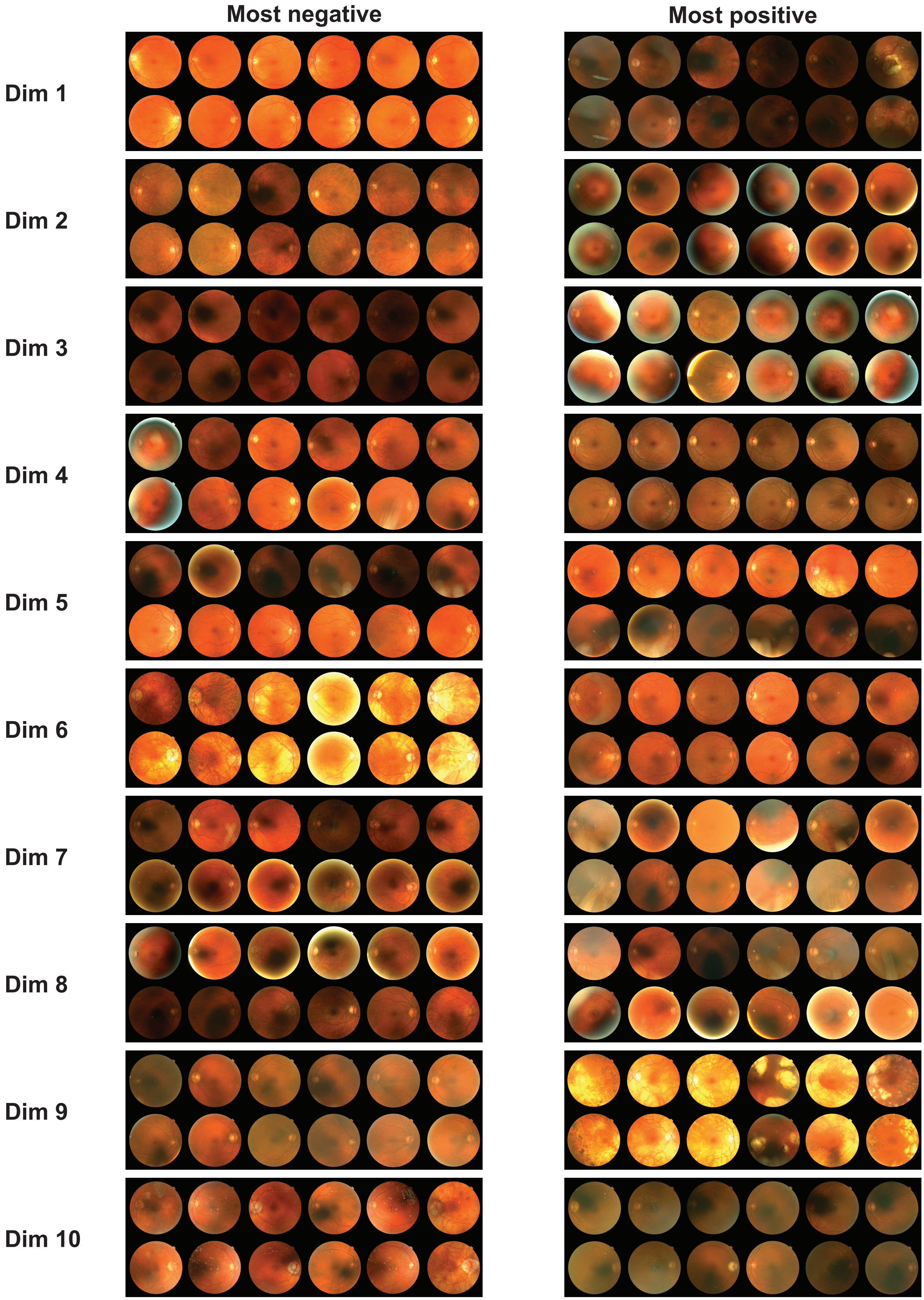


**Supplementary Figure 3. Illustration of information in the retinal fundus images captured by the feature dimensions, based on ImageNet pre-training.**

Shown are example images activating individual dimensions. For each dimension retinal fundus images are displayed for the left eye (upper row) and the right eye (lower row). Left column ("most negative") shows the individuals activating the corresponding feature dimension the most in negative direction; right column ("most positive") shows the individuals activating the corresponding feature dimension the most in positive direction. Reproduced by kind permission of UK Biobank ©.

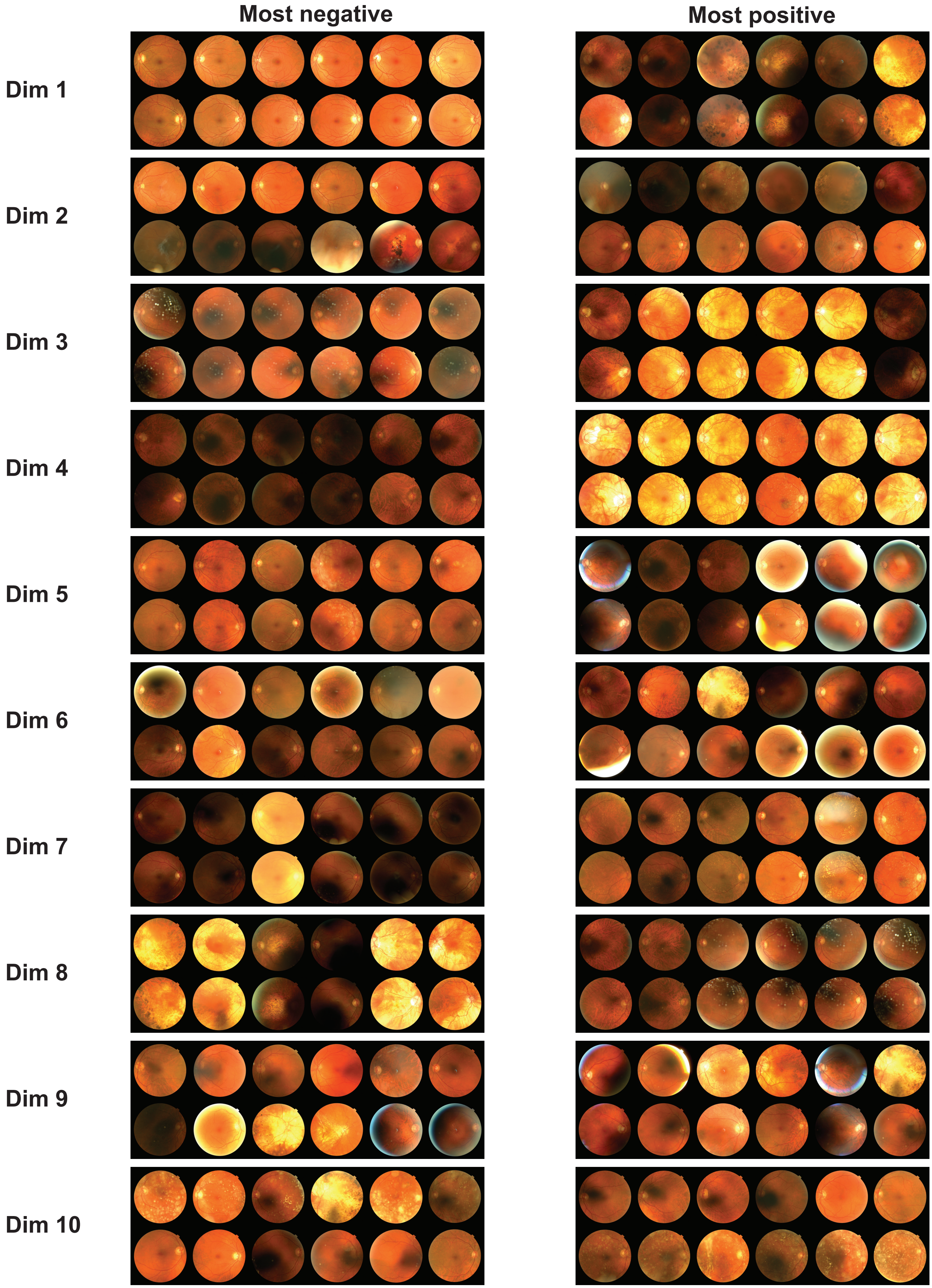


**Supplementary Figure 4. Illustration of information in the retinal fundus images captured by the feature dimensions, based on EyePACS pre-training.**
Shown are example images activating individual dimensions. For each dimension retinal fundus images are displayed for the left eye (upper row) and the right eye (lower row). Left column ("most negative") shows the individuals activating the corresponding feature dimension the most in negative direction; right column ("most positive") shows the individuals activating the corresponding feature dimension the most in positive direction. Reproduced by kind permission of UK Biobank ©.

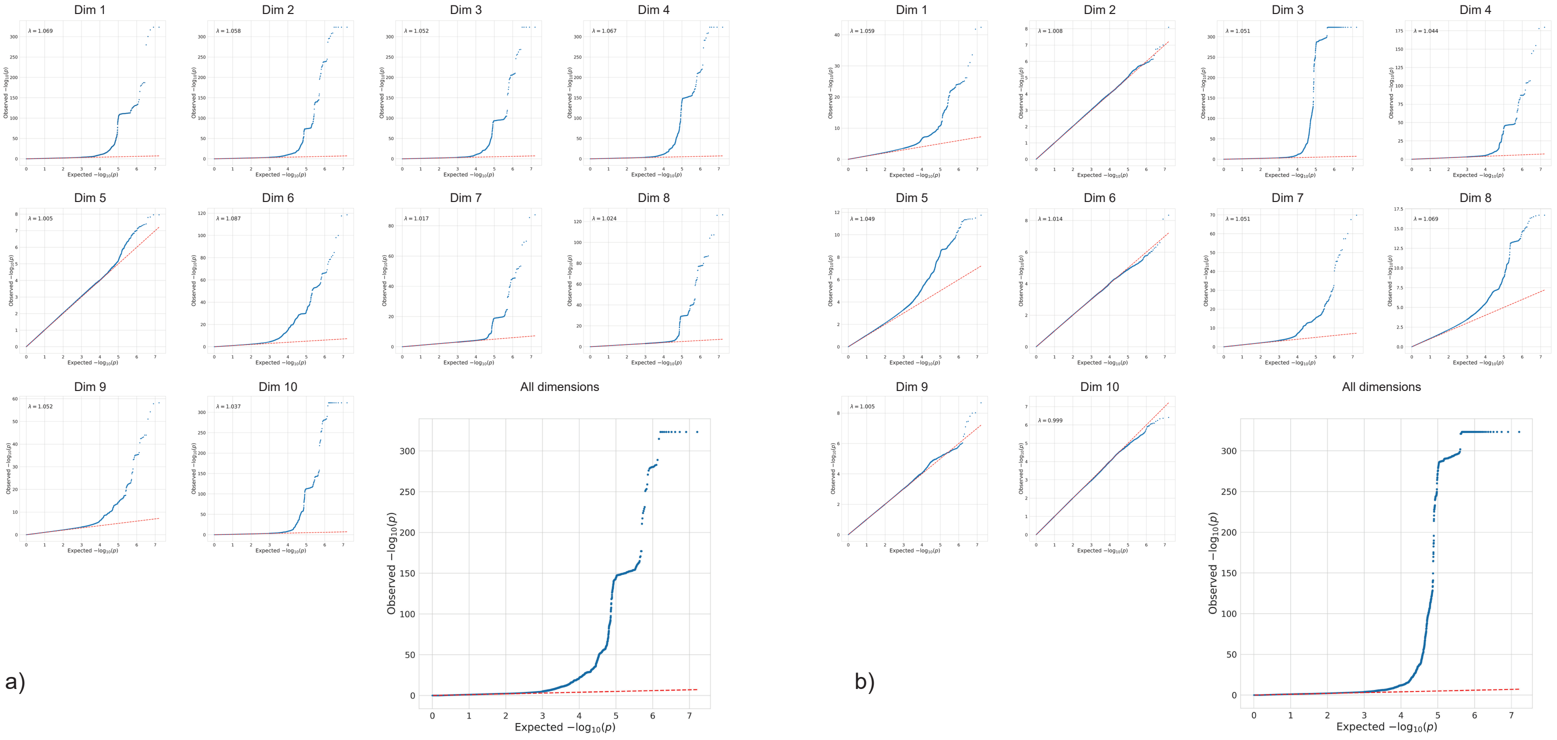


**Supplementary Figure 5. Distribution of the p-values from the GWAS based on ImageNet & EyePACS pre-training.** Quantile-quantile plot of observed p-values from the GWA study with ImageNet (a) and EyePACS (b) pre-training versus expected values. p-values are capped at $10^{-320}$.

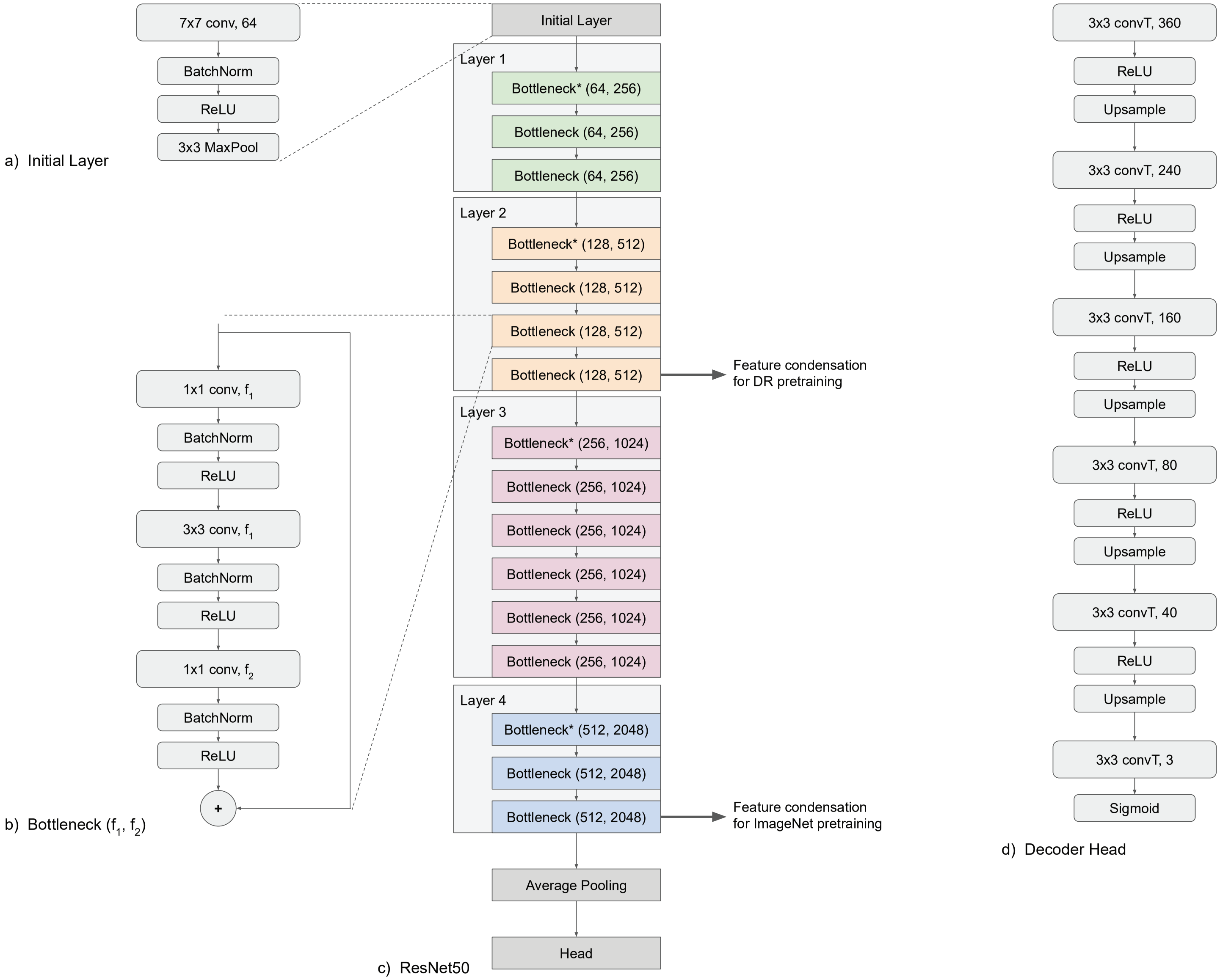


**Supplementary Figure 6. Architecture of the network used in the EyePACS pre-training.**
ResNet50 and Decoder architecture, as implemented in the pytorch library. Convolutional layers are depicted as blocks with “NxN conv, f”, where NxN is the kernel size, and f the number of convolutional filters. More details on skip connections and ResNet architecture can be found in the original paper.[35] Features are condensed as indicated, after the “1x1 conv, $f_2$” part of the corresponding Bottleneck block. a) Initial layer. b) One bottleneck block with $f_1$ and $f_2$ filters. Input and output of the block are connected via skip connection. c) ResNet50 architecture. Bottleneck* blocks are identical to Bottleneck blocks except for an additional downsampling step (not depicted). The head in the ImageNet pre-trained case is a simple linear layer. In the EyePACS pre-trained case the head splits into a decoder (see d)) for reconstructing the image and a linear layer for predicting the disease state. d) Decoder head for the image reconstruction for EyePACS pre-training. Reconstructed images are of size 224x224px instead of the input size of 448x448px. convT denotes transposed convolutions (or fractionally strided convolutions, sometimes also called deconvolutions) with no padding. Upsampling was always performed with a factor of 2 via bilinear interpolation.

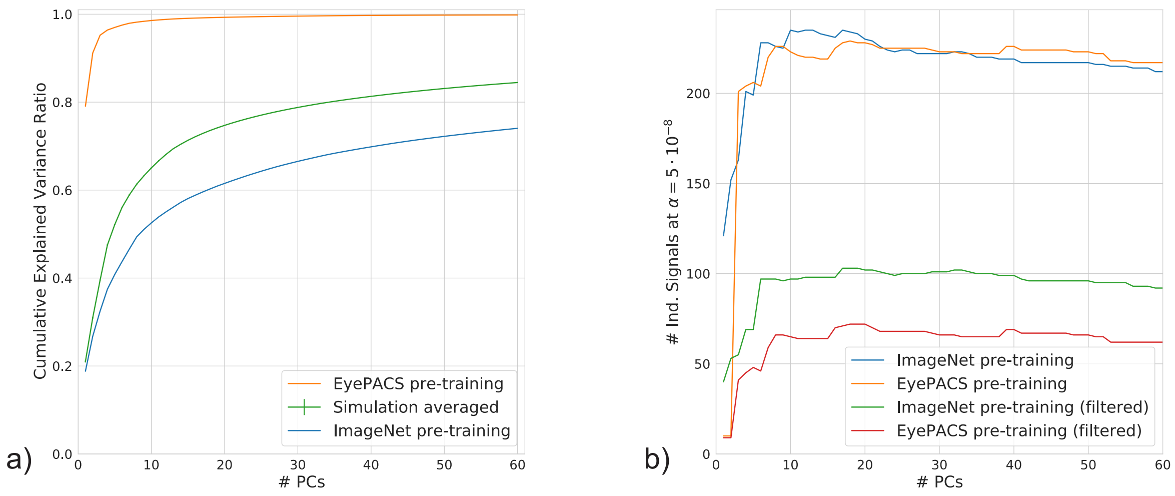


**Supplementary Figure 7. Effect of the number of feature dimensions/PCs.**
a) Relationship between the number of extracted principal components in transferGWAS and the explained variance of all feature dimensions. Shown is the cumulative explained variance ratio of the full deep representation by the first principal components, against the corresponding number of extracted principal components. Simulation results (with ImageNet pre-training) are averaged over ten runs. Standard error of the mean bars are shown but are very small due to very low differences between the simulations.
b) Relationship between the number of extracted principal components in transferGWAS and the number of independent genome-wide significant signals. Shown is the number of independent significant signals (as defined by PLINK clumping with window size of 250kb and $R^2$>0.1) as a function of the number of dimensions, for both the ImageNet pre-training and the EyePACS pre-training, with and without filtering variants with imputation quality scores <0.8 (imputation quality is always > 0.4) and MAF <1% (MAF is always >0.1%). P-values are always Bonferroni-corrected for the corresponding number of dimensions.

#### Supplementary Tables

**Supplementary Table 1:** Comparison of the results of association tests between eye-related traits and diseases and the feature dimensions obtained from the different pre-trainings.

| Eye-related trait | ImageNet Significant dimensions | p-value | EyePACS Significant dimensions | p-value | Cases |
|---|---|---|---|---|---|
| Myopia* | 1, 2, 3, 4, 5, 6', 7, 8, 9, 10 | $7.9 \cdot 10^{-296+}$ | 1, 3, 4, 5, 7, 8' | $7.7 \cdot 10^{-293+}$ | 30.6% |
| Hypermetropia* | 1, 2, 3, 4, 5, 6', 7, 8, 9, 10 | $1.4 \cdot 10^{-58+}$ | 3, 4, 5, 7, 8' | $3.7 \cdot 10^{-57+}$ | 14.8% |
| Presbyopia* | 1, 2, 4', 5, 6, 10 | $3.9 \cdot 10^{-67+}$ | 1', 3, 4, 5, 8 | $1.0 \cdot 10^{-63+}$ | 32.4% |
| Astigmatism* | 1, 2, 4', 6 | $1.9 \cdot 10^{-23+}$ | 1', 3, 8 | $4.2 \cdot 10^{-38+}$ | 10.0% |
| Strabismus* | 2, 3', 9 | $8.1 \cdot 10^{-15+}$ | 7, 8' | $5.3 \cdot 10^{-14+}$ | 1.4% |
| Amblyopia* | 2, 3', 7, 9 | $2.4 \cdot 10^{-20+}$ | 1, 7, 8' | $1.4 \cdot 10^{-16+}$ | 2.7% |
| Other eye condition* | | $2.4 \cdot 10^{-4}$ | 4' | $1.3 \cdot 10^{-6+}$ | 1.0% |
| Diabetes related eye disease** | | $8.9 \cdot 10^{-6}$ | 1' | $1.3 \cdot 10^{-7+}$ | 0.8% |
| Glaucoma** | | $7.3 \cdot 10^{-6}$ | 1' | $3.7 \cdot 10^{-7+}$ | 1.7% |
| Injury or trauma** | | $1.0 \cdot 10^{-2}$ | 4' | $7.0 \cdot 10^{-8+}$ | 0.5% |
| Cataract** | 1, 2, 4, 8, 10' | $2.0 \cdot 10^{-24+}$ | 1', 4 | $4.2 \cdot 10^{-44+}$ | 4.6% |
| Macular degeneration** | | $1.4 \cdot 10^{-5}$ | 4', 7 | $3.5 \cdot 10^{-27+}$ | 1.1% |
| Other serious eye condition** | 4' | $9.1 \cdot 10^{-11+}$ | 1', 4 | $9.9 \cdot 10^{-16+}$ | 1.8% |
| No eye disease** | 1,2, 4', 8, 10 | $2.0 \cdot 10^{-24+}$ | 1', 4 | $2.6 \cdot 10^{-67+}$ | 90.4% |

Shown are association testing results between specific eye-related traits and diseases and the 10 feature dimensions after ImageNet and EyePACS pre-training, respectively. For each trait, smallest p-values (Bonferroni-adjusted for 10 feature dimensions) and all significant feature dimension are shown.
*Traits are subcategories of the touchscreen question of reasons for wearing glasses/contact lenses. Available nonmissing values: $n$ = 46,140. The subcategory "Don't know" (7% of answers) was not analyzed in the PheWAS.
**Traits are subcategories of the touchscreen question of eye problems/disorders. Available nonmissing values: $n$ = 39,308.
$^{+}$Significant at Bonferroni-adjusted significance level $4.3 \cdot 10^{-6}$.
'Dimension with strongest association.

**Supplementary Table 2:** Empirical Power estimates from the simulation GWAS.

| Sample Size | Explained Variance of Latent Code | Empirical Power | SEM |
|---|---|---|---|
| 6,000 | 0.5 | 0.00% | 0.00% |
| 12,000 | 0.5 | 0.08% | 0.03% |
| 24,000 | 0.5 | 1.70% | 0.08% |
| 46,731 | 0.5 | 11.81% | 0.21% |
| 6,000 | 0.7 | 0.02% | 0.01% |
| 12,000 | 0.7 | 0.38% | 0.04% |
| 24,000 | 0.7 | 4.74% | 0.13% |
| 46,731 | 0.7 | 23.93% | 0.51% |

Shown are empirical estimates of the power in the simulation studies, defined as the fraction of discovered causal SNPs with p-value below the genome-wide significant threshold of $5 \cdot 10^{-8}$ relative to all causal SNPs, averaged over 10 runs, together with an estimate of the standard error of the mean (SEM). Number of causal SNPs is always set to 1,250.

**Supplementary Table 3:** Characteristics of the study population of n=46,731 Caucasian participants from the UK Biobank.

| | |
|---|---|
| Gender (% male) | 45.95% |
| Age mean (std) | 56.53 (7.95), $n = 46,731$ |
| BMI mean (std) | 27.15 (4.65), $n = 46,576$ |
| Systolic Blood Pressure mean (std) | 137.16 (18.26), $n = 45,688$ |
| Diastolic Blood Pressure mean (std) | 81.78 (10.00), $n = 45,688$ |
| Smoker | Current: 8.40%, Never: 56.82%, Previous: 34.78%, $n = 46,597$ |
| Eye disease* | 9.62%, $n = 39,308$ |
| Wears glasses | 89.10%, $n = 46,704$ |
| Hair color | Blonde: 11.39%, Red: 4.47%, Light brown: 41.78%, Dark brown: 38.59%, Black: 3.77%, $n = 46,175$ |
| Skin color | Very fair: 8.19%, Fair: 72.09%, Light olive: 18.04%, Dark olive: 1.38%, Brown: 0.29%, $n = 46,288$ |
| Tanning | Get very tanned: 19.53%, Get moderately tanned: 40.41%, Get mildly or occasionally tanned: 22.19%, Never tan, only burn: 17.87%, $n = 45,735$ |

Shown are relative frequencies, relative to the number of nonmissing samples for the respective variable, or mean (standard deviation), as well as the number of available samples with nonmissing values. Missing values on touchscreen questions were due either to a selection of "Prefer not to answer" or "Do not know".
* Eye diseases included in this category are diabetes-related eye disease, glaucoma, injury or trauma resulting in loss of vision, cataract, macular degeneration, and other serious eye conditions.

**Supplementary Table 4:** Conditional Analysis for genomic region chr15:28.800-29.589Mb

| Variant | EAF | Info | Uncond. p-value | cSNP1 rs56166703 $R^2$ | cSNP1 rs56166703 P | cSNP2 rs72454513 $R^2$ | cSNP2 rs72454513 P | cSNP3 rs2670992 $R^2$ | cSNP3 rs2670992 P |
|---|---|---|---|---|---|---|---|---|---|
| rs56166703 | 0.53 | 0.93 | $\mathbf{5.5 \cdot 10^{-36}}$ | - | - | - | - | - | - |
| rs72454513* | 0.22 | 0.90 | $\mathbf{4.5 \cdot 10^{-27}}$ | 0.0583 | $\mathbf{5.8 \cdot 10^{-14}}$ | - | - | - | - |
| rs35615406 | 0.21 | 0.82 | $\mathbf{2.5 \cdot 10^{-10}}$ | 0.0004 | $4.2 \cdot 10^{-7}$ | 0.0167 | $1.6 \cdot 10^{-5}$ | 0.0001 | - |
| rs2670992+ | 0.61 | 0.87 | $\mathbf{2.0 \cdot 10^{-9}}$ | 0.7860 | $\mathbf{1.4 \cdot 10^{-12}}$ | 0.0355 | $\mathbf{1.1 \cdot 10^{-12}}$ | - | - |
| rs11263729′ | 0.12 | 0.84 | $1.1 \cdot 10^{-2}$ | 0.0891 | $\mathbf{1.7 \cdot 10^{-12}}$ | 0.0117 | $\mathbf{1.1 \cdot 10^{-9}}$ | 0.0600 | $1.8 \cdot 10^{-6}$ |

Shown are association results of the stepwise conditional analysis for the genomic region chr15:28.800-29.589Mb with EyePACS pre-training. EAF is effect allele frequency, Info is the imputation quality score of the variant. $R^2$ between variants and cSNPs are computed on study data
* Strongest remaining association after first round of adjustment.
+ Strongest remaining association after second round of adjustment.
′ Strongest remaining association after second round of adjustment.

**Supplementary Table 5:** Number of independent significant loci shared between or unique to *transferGWAS* and baseline GWAS with standard traits (only on non-imputed data).

| Baseline traits | ImageNet both | ImageNet only *transferGWAS* | ImageNet only baseline | EyePACS both | EyePACS only *transferGWAS* | EyePACS only baseline |
|---|---|---|---|---|---|---|
| Eye-traits | 3 | 71 | 125 | 2 | 24 | 126 |
| Eye & physical traits | 3 | 71 | 218 | 3 | 23 | 218 |
| Eye, physical, & health | 3 | 71 | 220 | 3 | 23 | 220 |

Shown are the number of significant loci discovered either only by baseline GWAS on standard phenotypical traits from the UKB (baseline traits) or only in *transferGWAS*. "both" denotes loci that were found both by the baseline GWAS and the *transferGWAS* (mapped to ±250kb).

Jorge Cuadros and George Bresnick. Eyepacs: an adaptable telemedicine system for diabetic retinopathy screening. *Journal of diabetes science and technology*, 3(3):509–516, 2009.

Huazhu Fu, Boyang Wang, Jianbing Shen, Shanshan Cui, Yanwu Xu, Jiang Liu, and Ling Shao. Evaluation of retinal image quality assessment networks in different color-spaces. In *International Conference on Medical Image Computing and Computer-Assisted Intervention*, pages 48–56. Springer, 2019.

**Supplementary Table 6:** Empirical power of simulated baseline GWAS results for 1 and 10 simulated standard traits.

| | **20% explained variance** | | **50% explained variance** | |
|---|---|---|---|---|
| | **1 trait** | **10 traits** | **1 trait** | **10 traits** |
| 50% sparsity | 1.4% (±0.16%) | 9.0% (±0.39%) | 8.9% (±0.29%) | 46.2% (±0.85%) |
| 80% sparsity | 2.1% (±0.08%) | 12.3% (±0.13%) | 9.1% (±0.25%) | 50.5% (±0.56%) |
| 95% sparsity | 3.2% (±0.15%) | 20.6% (±0.26%) | 8.1% (±0.15%) | 52.7% (±0.63%) |

GWAS results for 1 and 10 simulated standard traits. *transferGWAS* achieves empirical power of 15.5% (±0.40%). Power is averaged over three random seeds. Numbers in parentheses denote the SEM.


Po-Ru Loh, George Tucker, Brendan K Bulik-Sullivan, Bjarni J Vilhjalmsson, Hilary K Finucane, Rany M Salem, Daniel I Chasman, Paul M Ridker, Benjamin M Neale, Bonnie Berger, et al. Efficient bayesian mixed-model analysis increases association power in large cohorts. *Nature genetics*, 47(3):284–290, 2015.

Po-Ru Loh, Gleb Kichaev, Steven Gazal, Armin P Schoech, and Alkes L Price. Mixed-model association for biobank-scale datasets. *Nature genetics*, 50(7):906–908, 2018.

Mitchell J Machiela and Stephen J Chanock. Ldlink: a web-based application for exploring population-specific haplotype structure and linking correlated alleles of possible functional variants. *Bioinformatics*, 31(21):3555–3557, 2015.

Louise AC Millard, Neil M Davies, Tom R Gaunt, George Davey Smith, and Kate Tilling. Software application profile: Phesant: a tool for performing automated phenome scans in uk biobank. *International journal of epidemiology*, 2017.

Olga Russakovsky, Jia Deng, Hao Su, Jonathan Krause, Sanjeev Satheesh, Sean Ma, Zhiheng Huang, Andrej Karpathy, Aditya Khosla, Michael Bernstein, et al. Imagenet large scale visual recognition challenge. *International journal of computer vision*, 115(3):211–252, 2015.

Milly S Tedja, Robert Wojciechowski, Pirro G Hysi, Nicholas Eriksson, Nicholas A Furlotte, Virginie JM Verhoeven, Adriana I Iglesias, Magda A Meester-Smoor, Stuart W Tompson, Qiao Fan, et al. Genome-wide association meta-analysis highlights light-induced signaling as a driver for refractive error. *Nature genetics*, 50(6):834–848, 2018.
