## Supplementary File 3 - Regional Plots for "TransferGWAS: GWAS of images using deep transfer learning"

| Locus 1: chr1:23.182-23.682Mb - **ImageNet only** (rs1208930/1:23432346_C/T) | 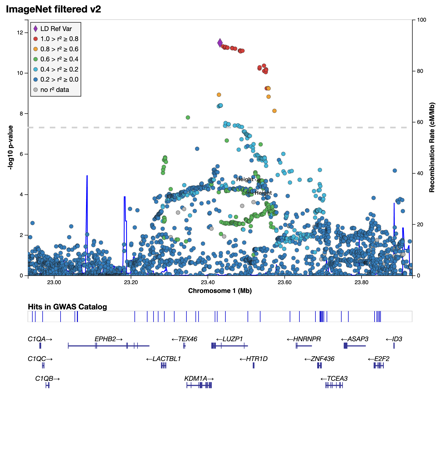 | 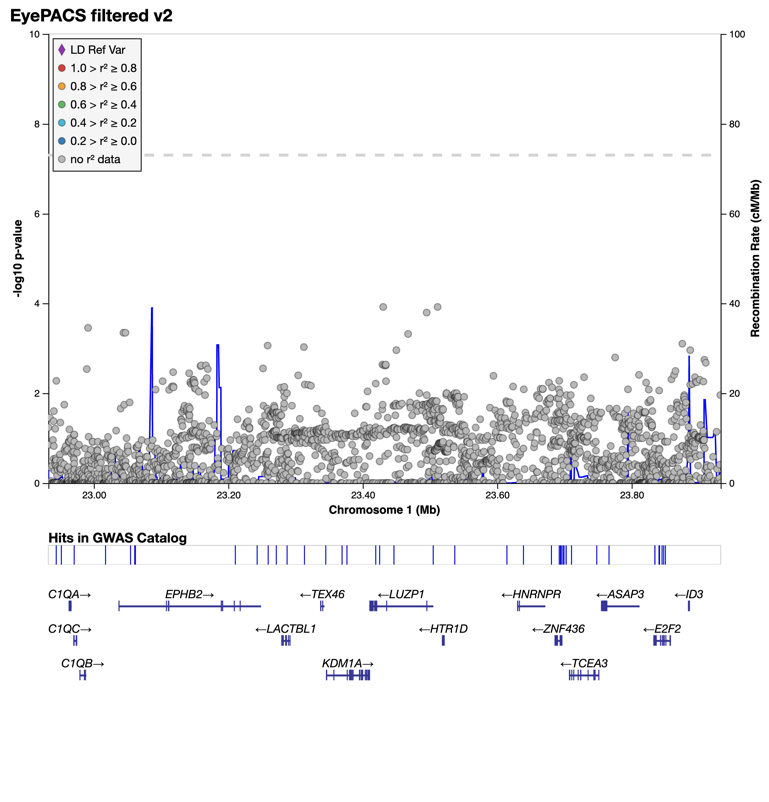 |
| --- | --- | --- |
| Locus 2: chr1:91.921-92.421Mb - **ImageNet only** (rs284874/1:92170644_T/C) | 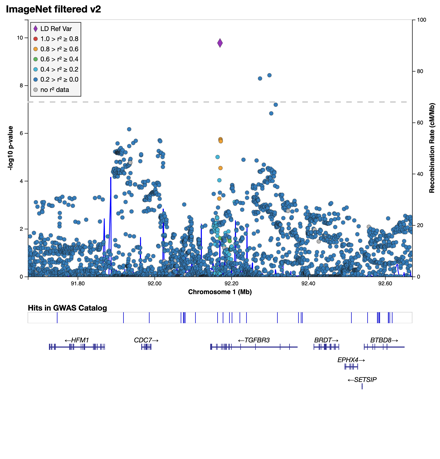 | 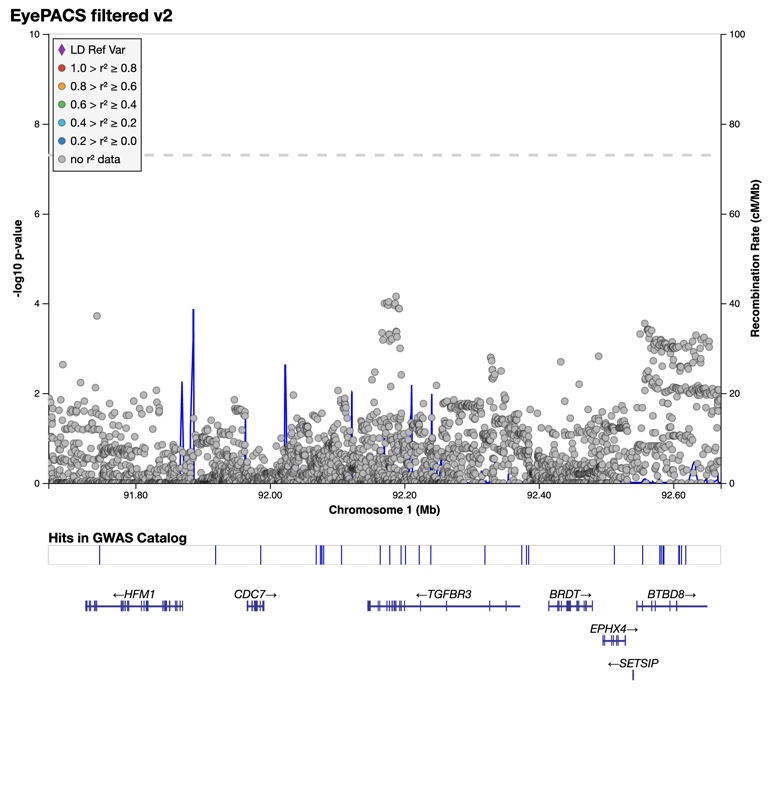 |
| Locus 3: chr1:110.382-110.882Mb - **ImageNet only** (rs12143283/1:110631977_A/C) | 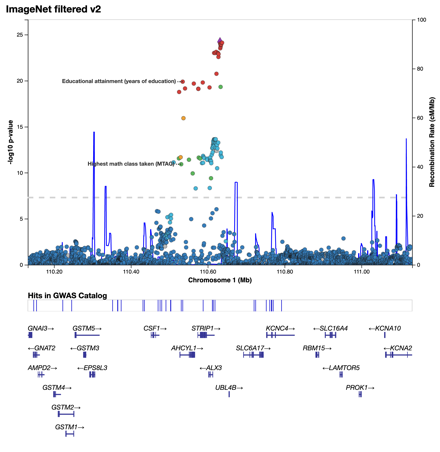 | 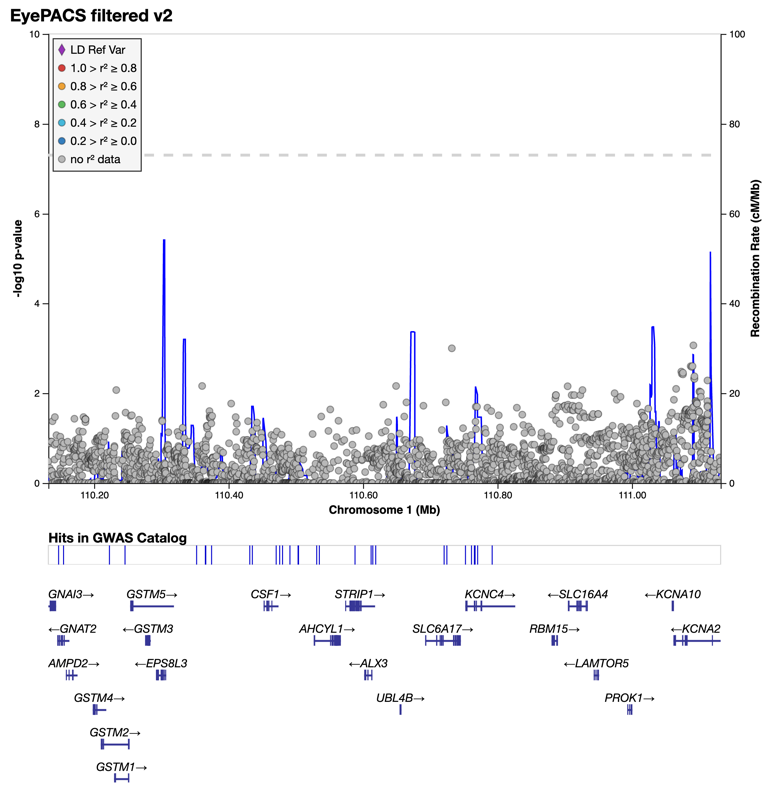 |
| Locus 4: chr1:149.999-150.758Mb - **ImageNet only** (rs35787821/1:150249349_G/GA) | 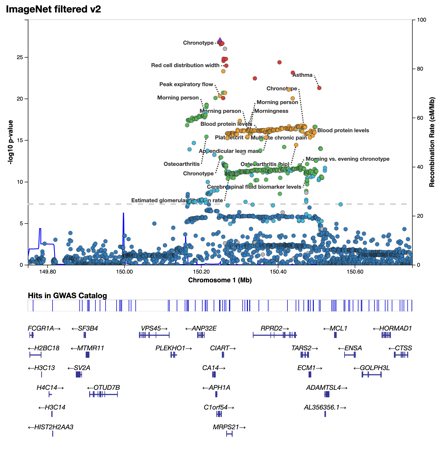 | 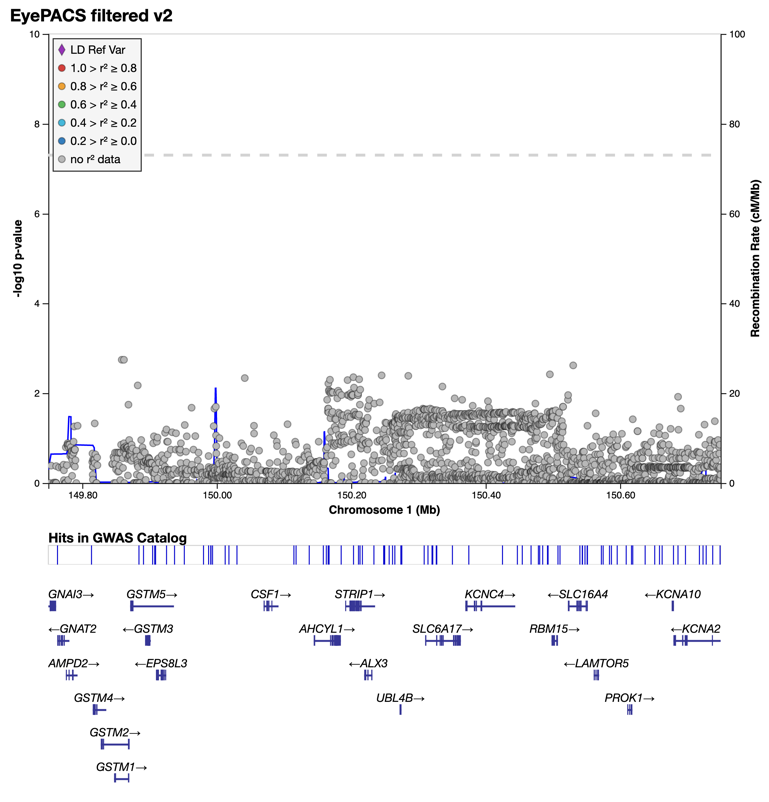 |
| Locus 4: chr1:149.999-150.758Mb - **ImageNet only** (rs71580314/1:150507691_A/AGGG) | 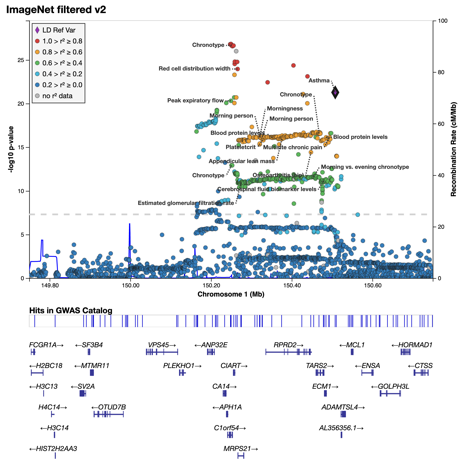 |  |
| Locus 5: chr1:204.865-205.432Mb - **ImageNet & EyePACS** (rs12048743/1:205114873_C/G) | 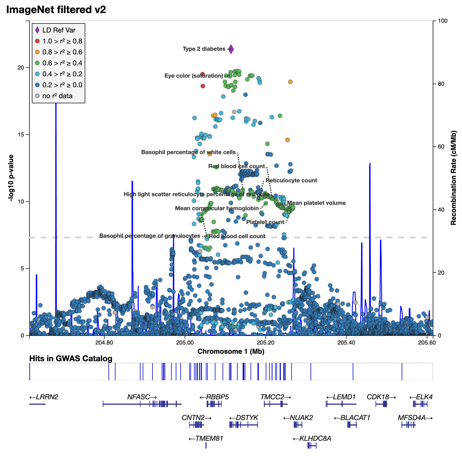 | 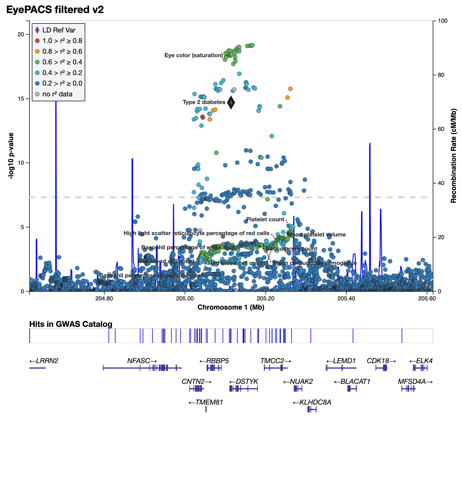 |
| Locus 5: chr1:204.865-205.432Mb - (1:205167799_C/G) | 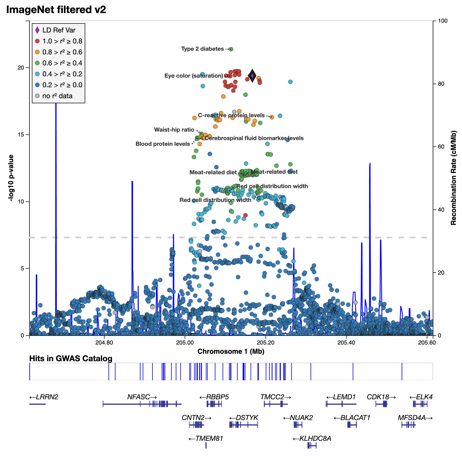 | 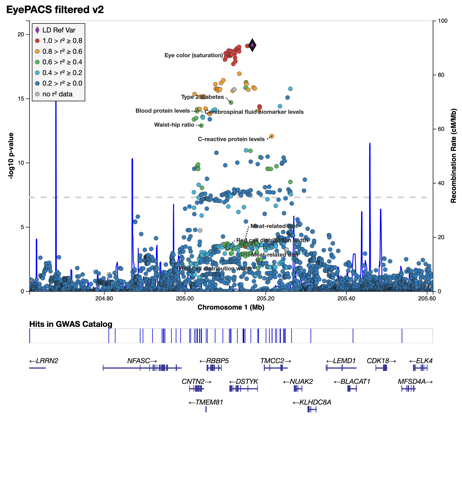 |
| Locus 5: chr1:204.865-205.432Mb - **ImageNet only** (rs6659069/1:205182031_A/G) | 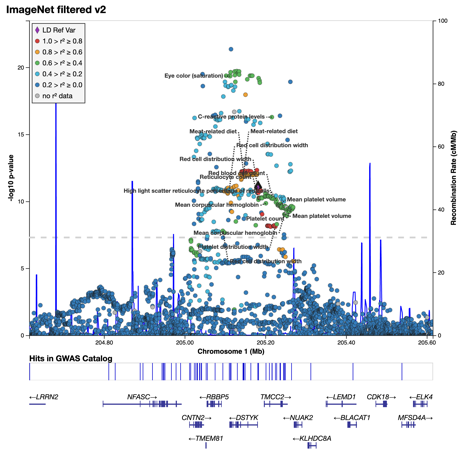 | Not Sign in EyePacs (P=0.01) |
| Locus 6: chr1:212.181-212.681Mb - **EyePACS, ImageNet subsign** (rs12747025/1:212430815_C/T) | 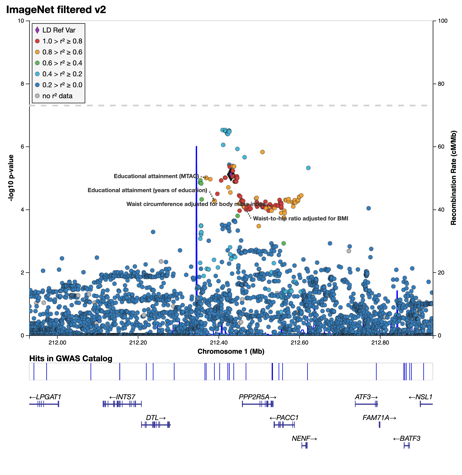 | 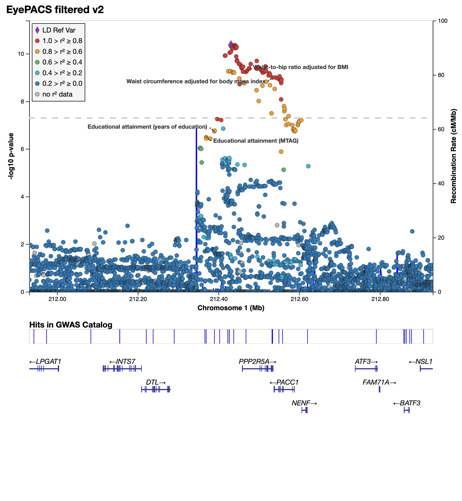 |
| Locus 7: chr1:226.81-227.31Mb - **ImageNet, EyePACS subsign** (rs1295646/1:227059739_G/C) | 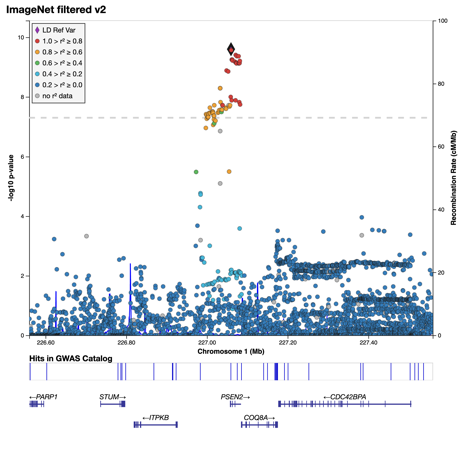 | 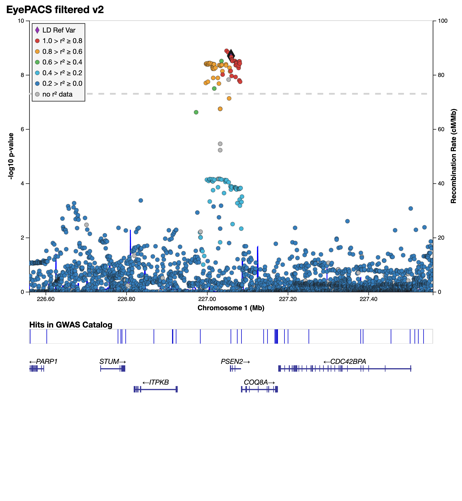 |
| Locus 8: chr2:0.014-0.514Mb  **ImageNet & EyePACS** (rs7595075/2:264019_C/A) | 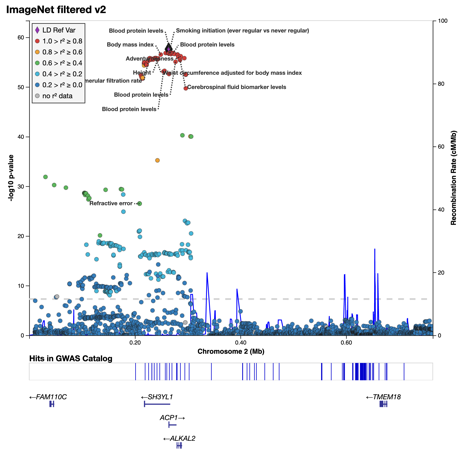 | 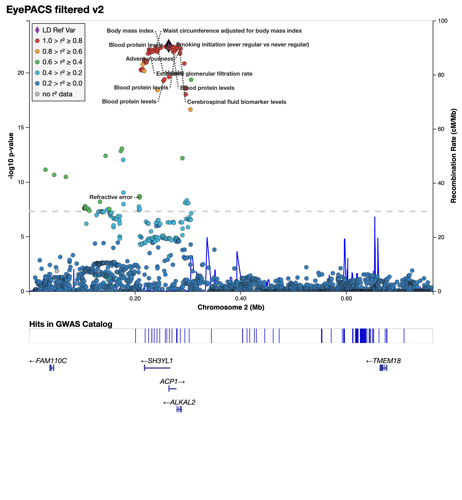 |
| Locus 9: chr2:134.03-134.53Mb - **ImageNet only** (rs1812081/2:134279837_G/T) | 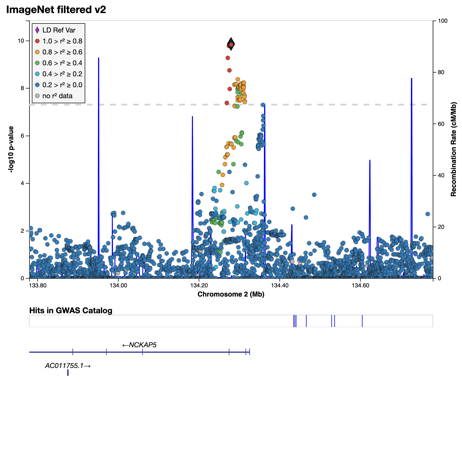 | 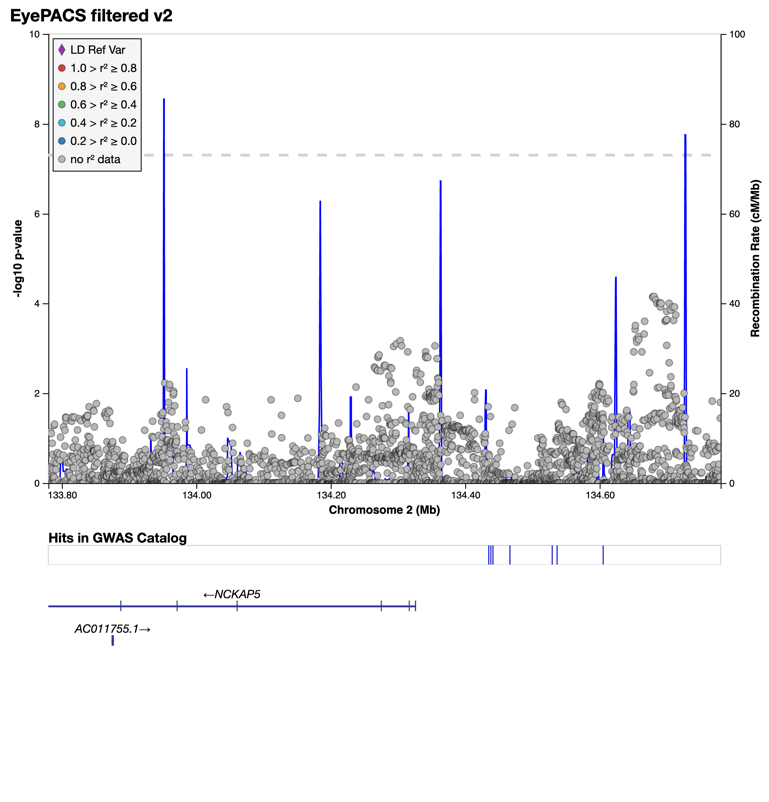 |
| Locus 10: chr2:218.421-218.921Mb - **ImageNet only** (rs2571461/2:218670945_T/G) | 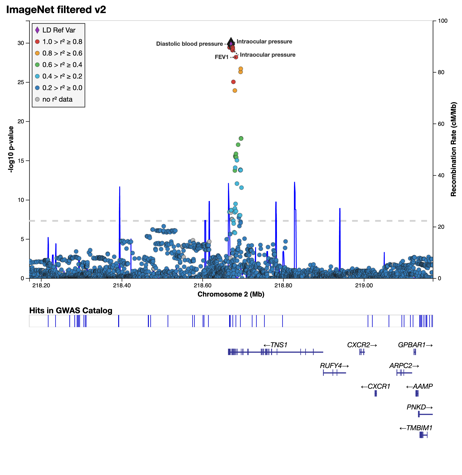 | 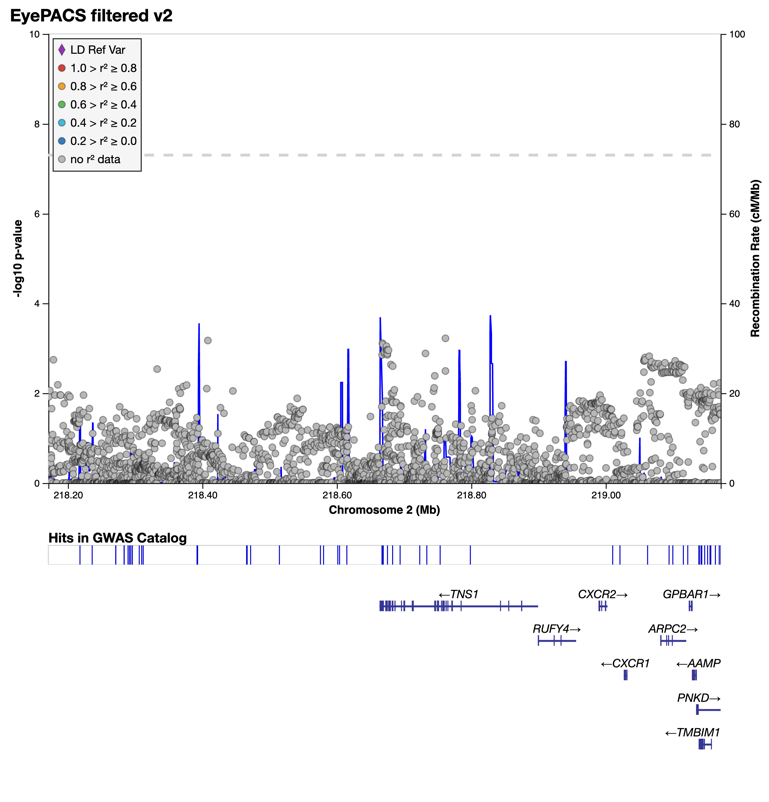 |
| Locus 11: chr3:4.152-4.652Mb  **EyePACS, ImageNet subsign**  (rs9837864 / 3:4402089_G/A) | 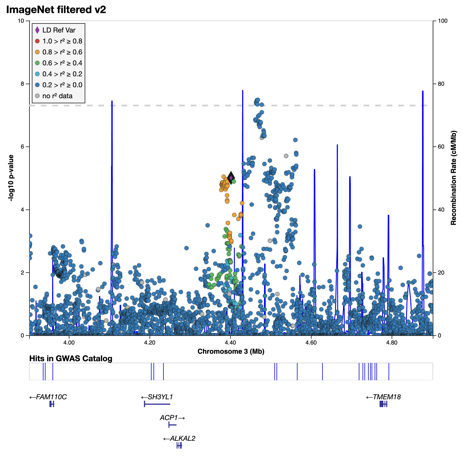 | 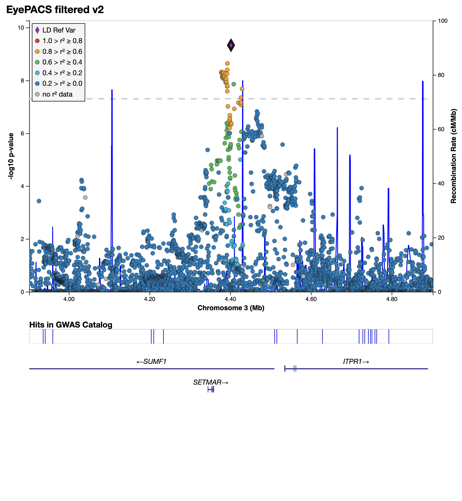 |
| Locus 11: chr3:4.152-4.652Mb  (rs9837864 / 3:4468598_C/T) | 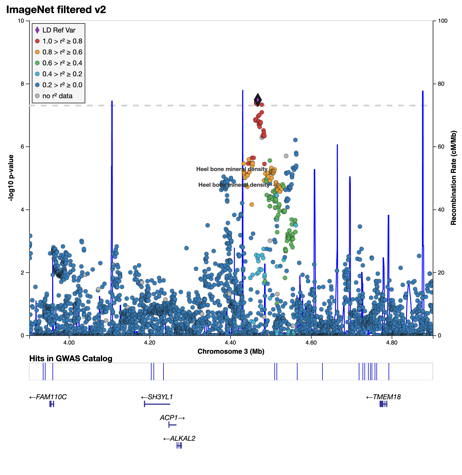 | 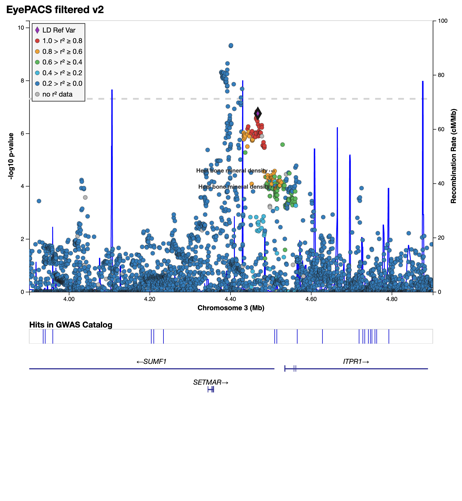 |
| Locus 12: chr3:41.031-41.531Mb  **EyePACS, ImageNet subsign** (rs4135387/3:41281413_T/A) | 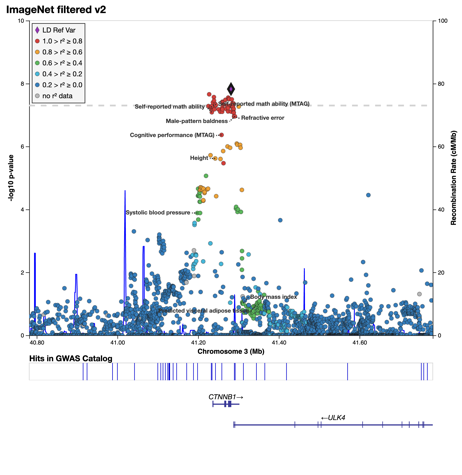 | 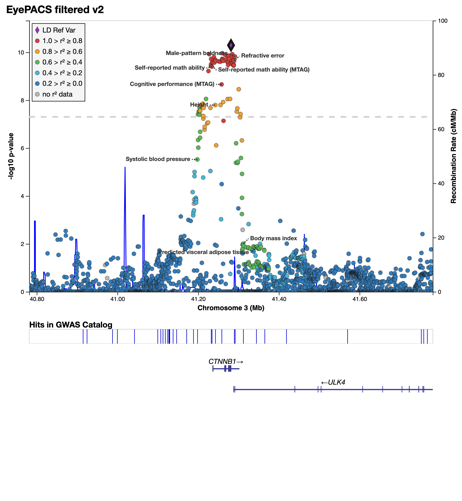 |
| Locus 13: chr4:84.781-85.281Mb  **ImageNet, EyePACS subsign** (rs2903945/4:85031002_A/T) | 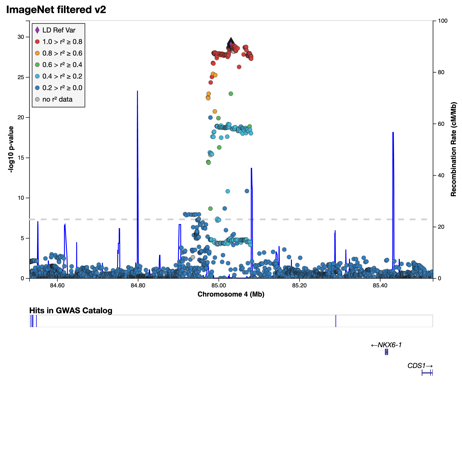 | 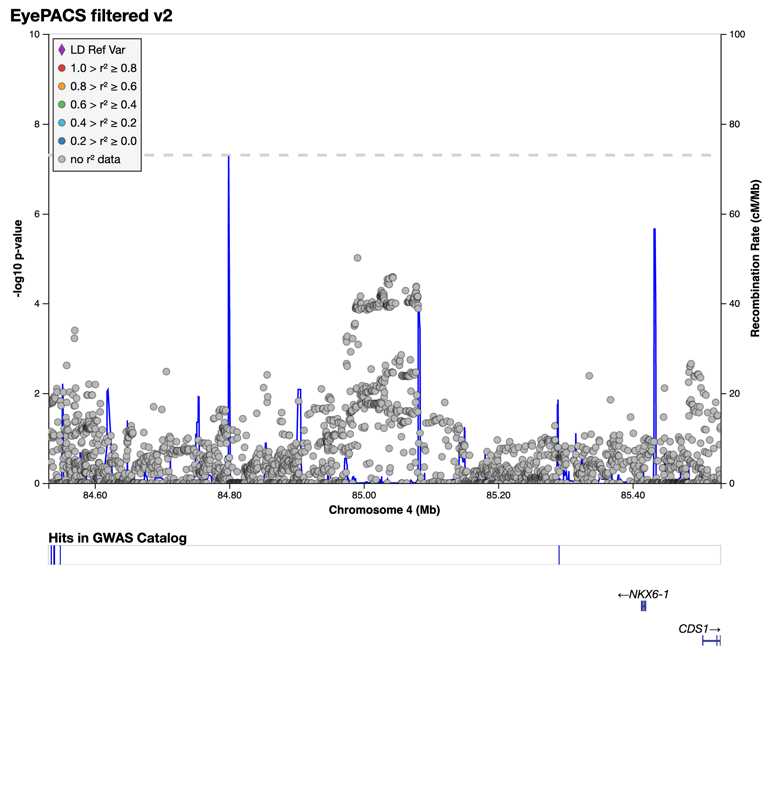 |
| Locus 14: chr4:119.736-120.807Mb - **ImageNet only** (rs12647722/4:119986395_G/T) | 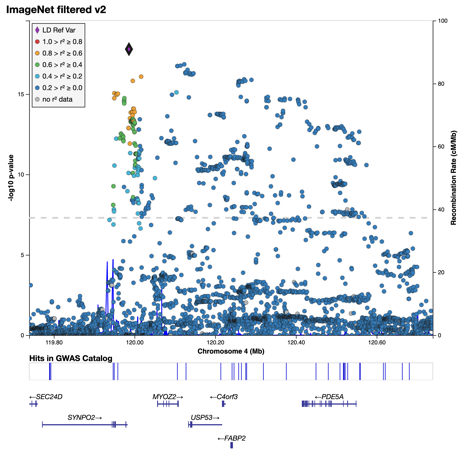 | 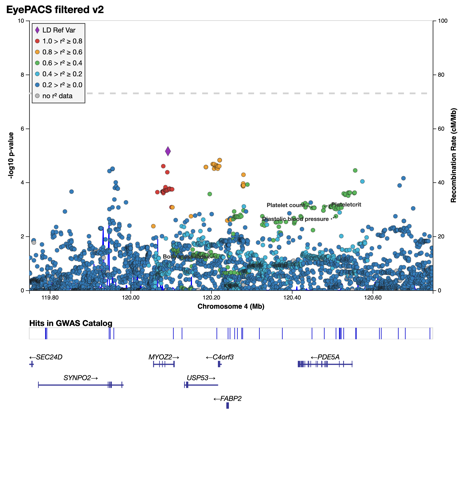  Overview of whole region, Ref SNP is SNP with smallest P-Value |
| Locus 14: chr4:119.736-120.807Mb - **ImageNet only** (rs141285286/4:120134796_G/GATATAT) | 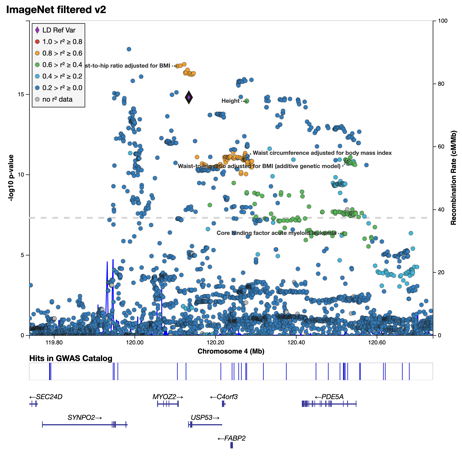 |  |
| Locus 14: chr4:119.736-120.807Mb - **ImageNet only** (rs10003742/4:120271798_G/C) | 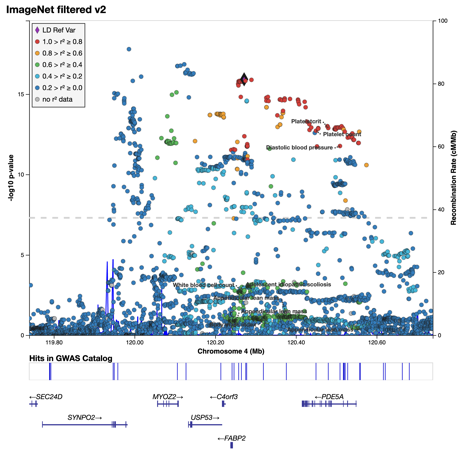 |  |
| Locus 14: chr4:119.736-120.807Mb - **ImageNet, EyePACS subsign** (rs2389875/4:120557014_G/T) | 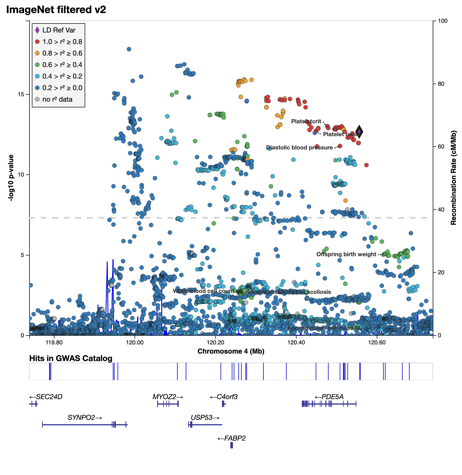 | 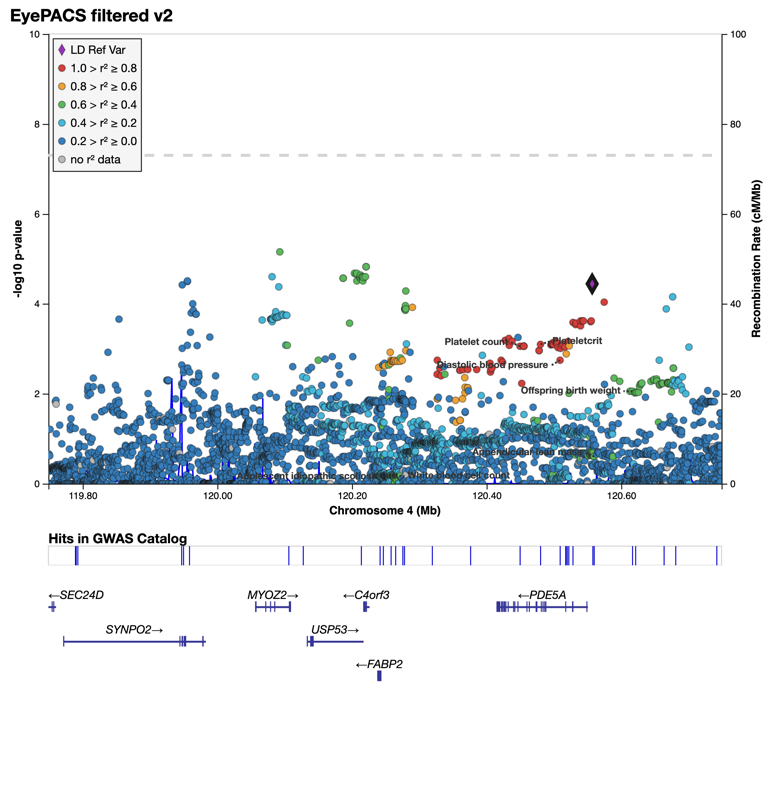 |
| Locus 15: chr5:33.674-34.202Mb - **ImageNet, EyePACS subsign** (rs982823/5:33923911_G/C) | 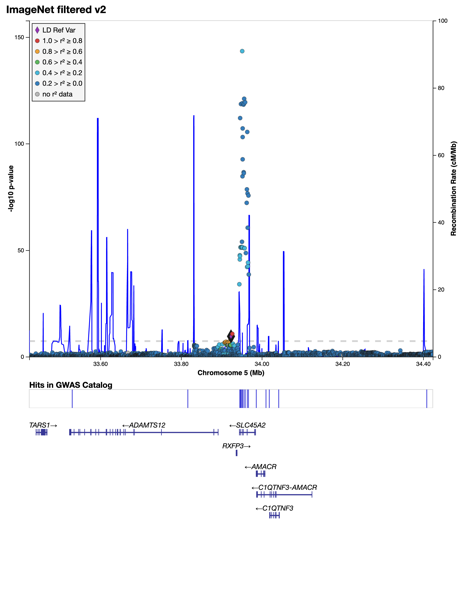 | 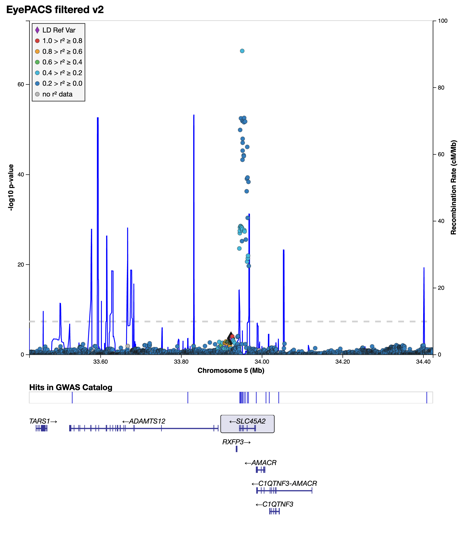  EyePacs P= 0.00028 |
| Locus 15: chr5:33.674-34.202Mb - **ImageNet & EyePACS** (rs16891982/5:33951693_C/G) | 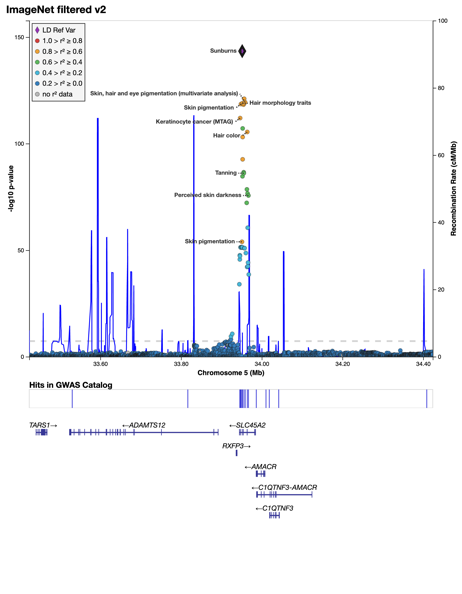 | 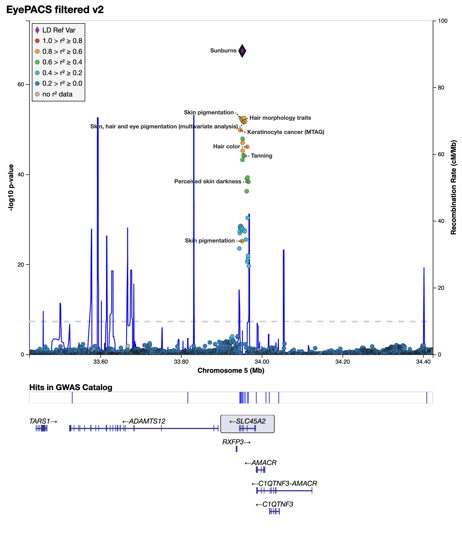 |
| Locus 16: chr5:77.656-78.238Mb - **ImageNet only** (5:77905667_TA_T/5:77905667_TA/T) | 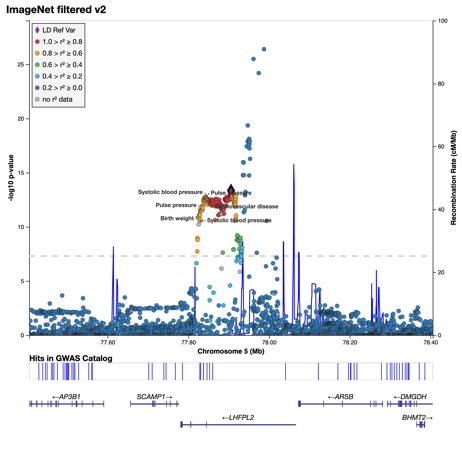 | 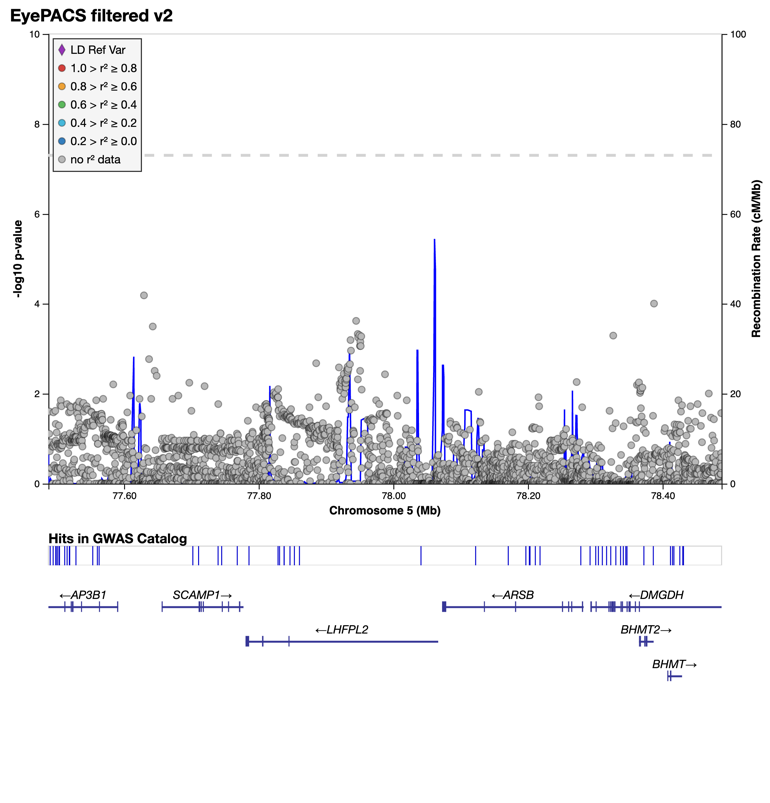 |
| Locus 16: chr5:77.656-78.238Mb - **ImageNet only** (5:77947233_CAAA_C/5:77947233_CAAA/C) | 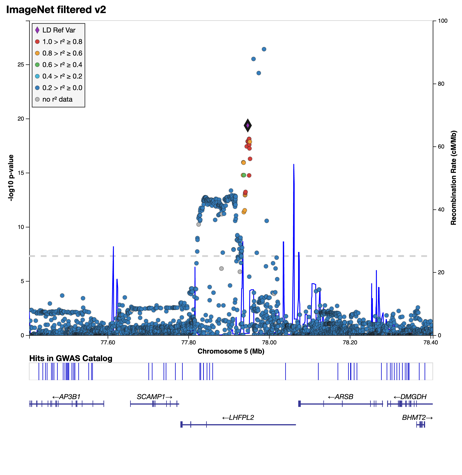 | 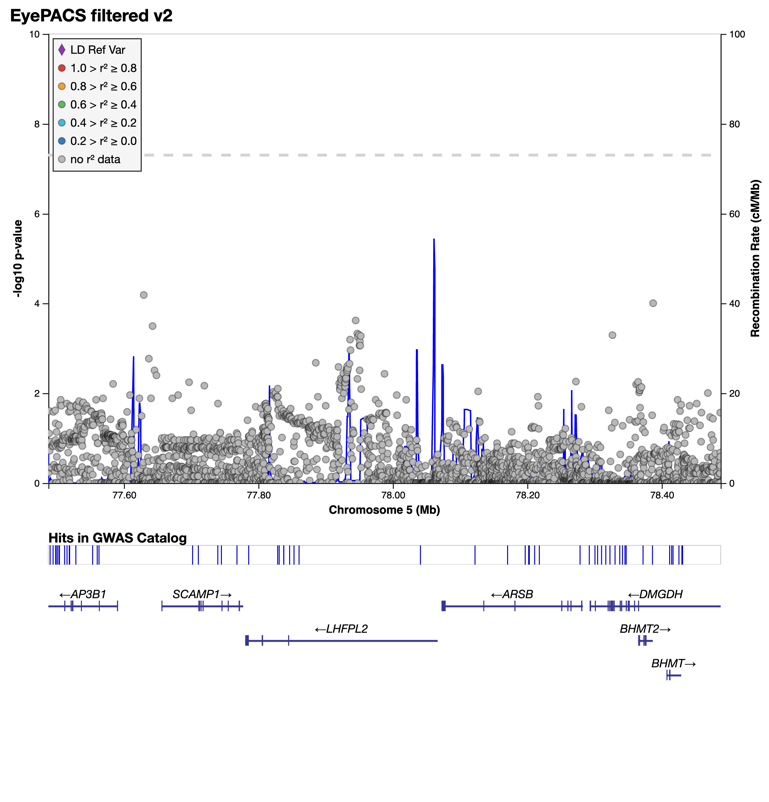 |
| Locus 16: chr5:77.656-78.238Mb - **ImageNet only** (rs784420/5:77987524_A/G) | 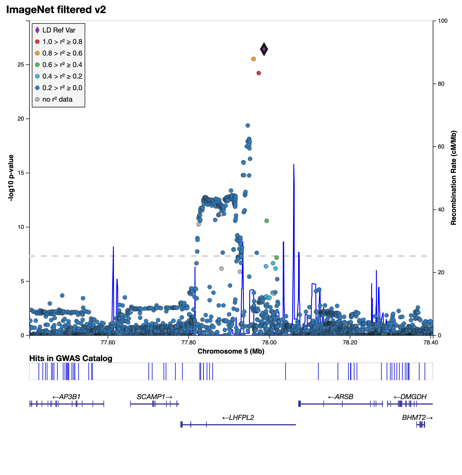 |  |
| Locus 17: chr5:87.526-88.096Mb - **EyePACS only** (rs199502002/5:87775507_T/TG) | Not Sign in ImageNet  (P= 0.0027) | 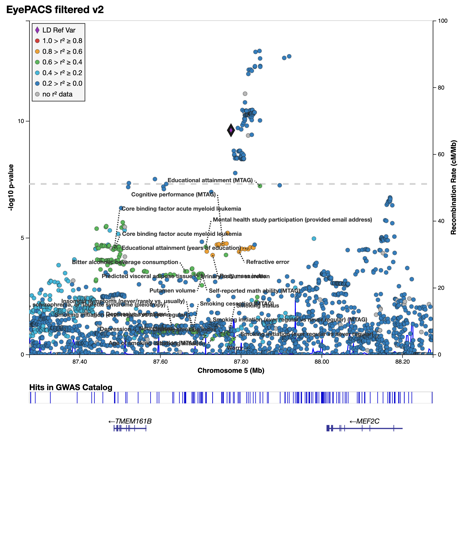 |
| Locus 17: chr5:87.526-88.096Mb - **EyePACS, ImageNet subsign** (5:87846165_GA_G/5:87846165_GA/G) | 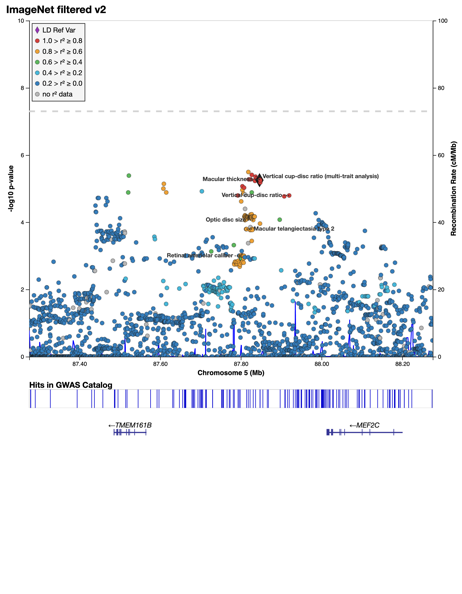 | 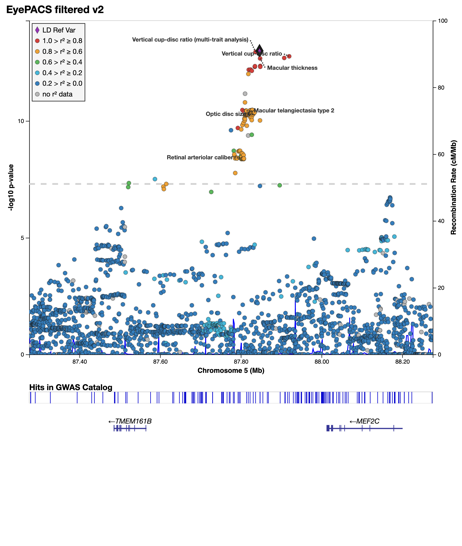 |
| Locus 18: chr6:0.146-0.866Mb  **ImageNet & EyePACS**  (rs12203592/6:396321_C/T) | 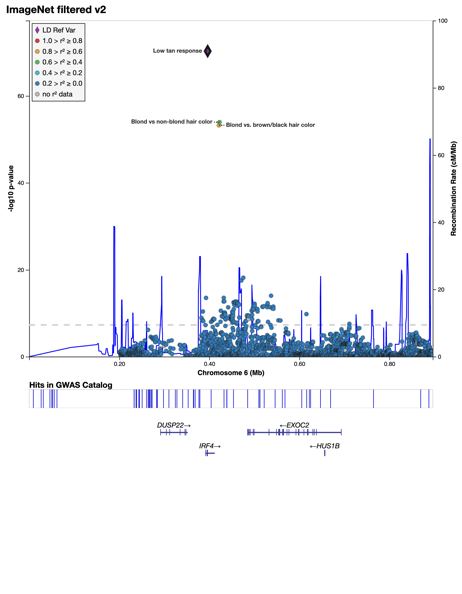 | 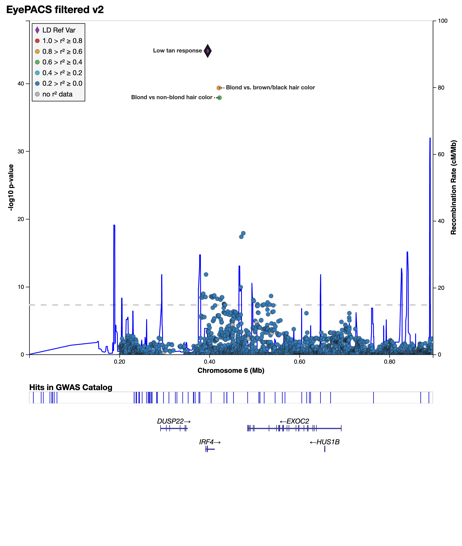 |
| Locus 18: chr6:0.146-0.866Mb - **ImageNet, EyePACS subsign** (rs6925797/6:432050_C/T) | 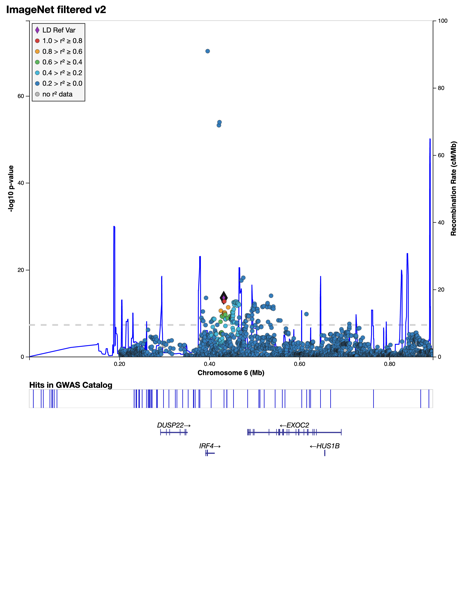 | 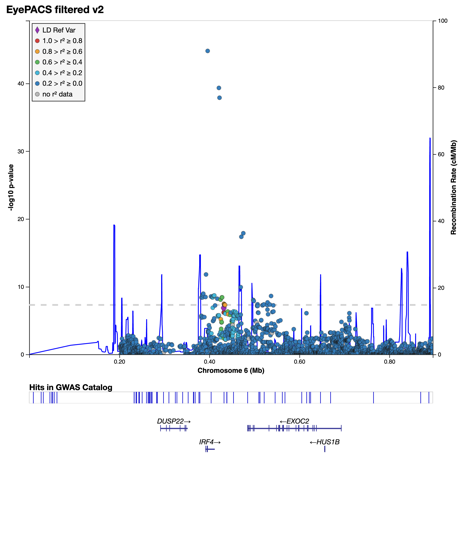 |
| Locus 18: chr6:0.146-0.866Mb - **ImageNet, EyePACS subsign** (rs1540767/6:466368_G/A) | 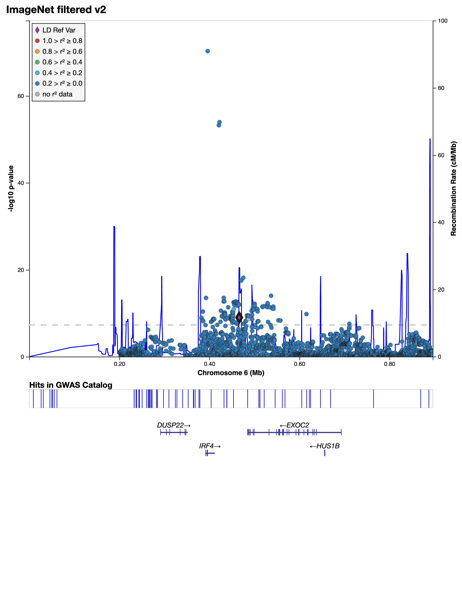 | 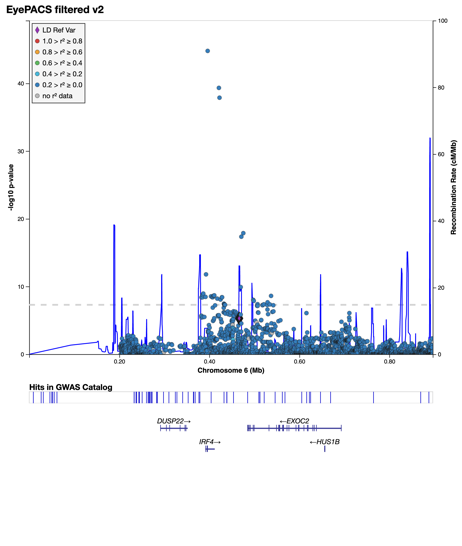 |
| Locus 18: chr6:0.146-0.866Mb  **ImageNet only**  rs950039/6:493976_G/A) | 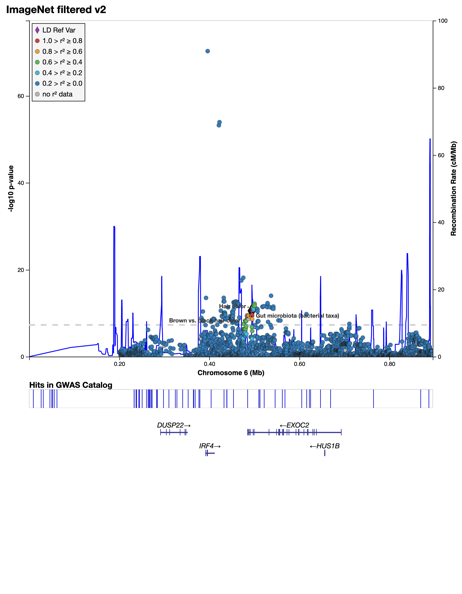 | EyePACS pval 0.0035 |
| Locus 18: chr6:0.146-0.866Mb - **ImageNet, EyePACS subsign** (rs11242982/6:537157_C/T) | 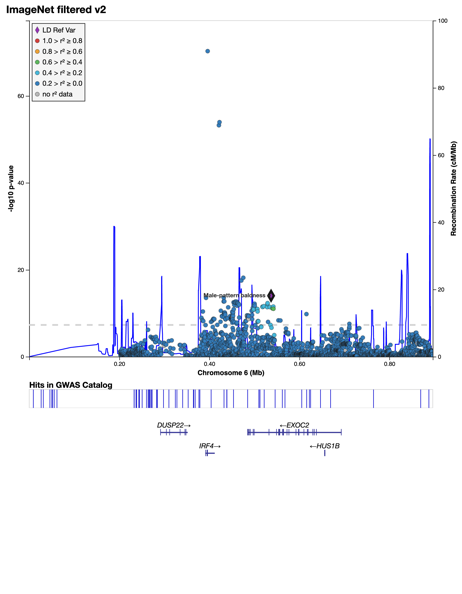 | 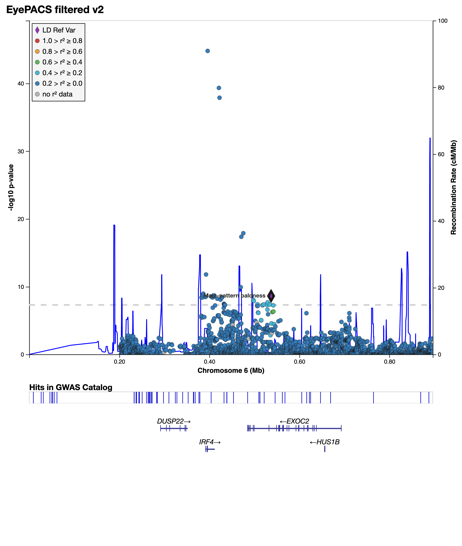 |
| Locus 18: chr6:0.146-0.866Mb - **ImageNet, EyePACS subsign** (rs74445037/6:615736_A/G) | 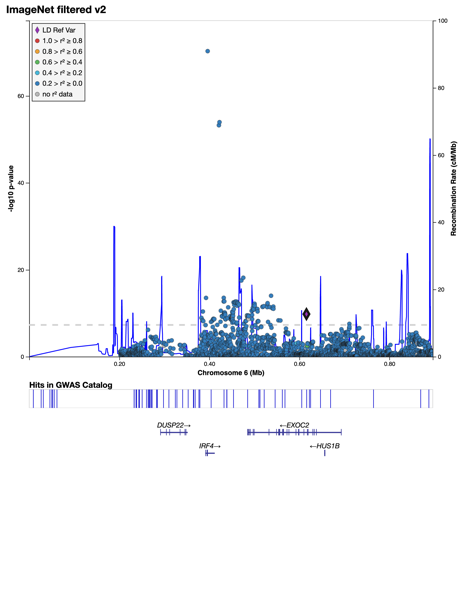 | 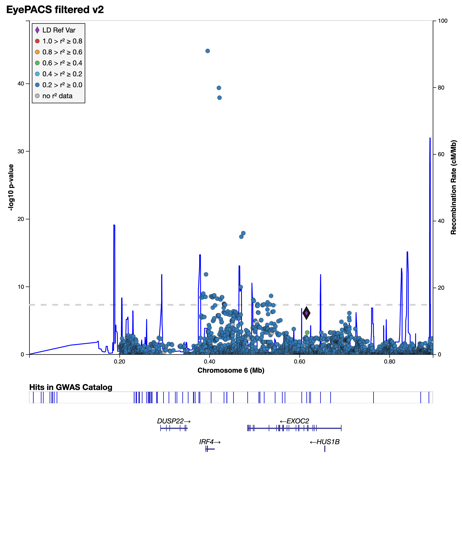 |
| Locus 19: chr6:1.136-1.636Mb  **EyePACS, ImageNet subsign** (rs9405472/6:1385814_C/A) | 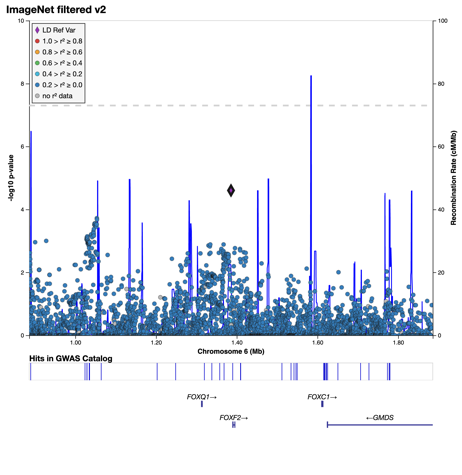 | 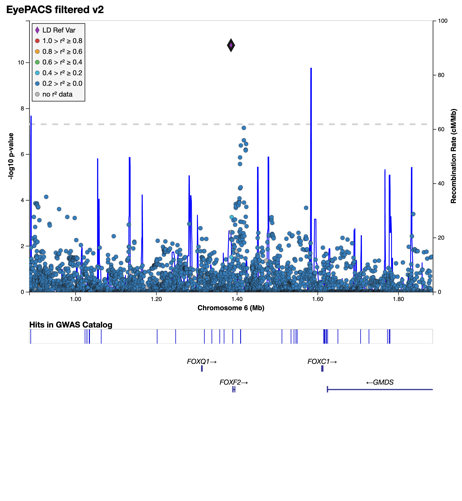 |
| Locus 20: chr6:6.656-7.156Mb - **ImageNet, EyePACS subsign** (rs9502516/6:6905830_G/A) | 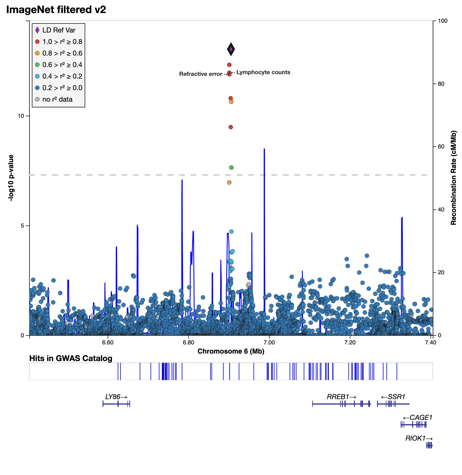 | 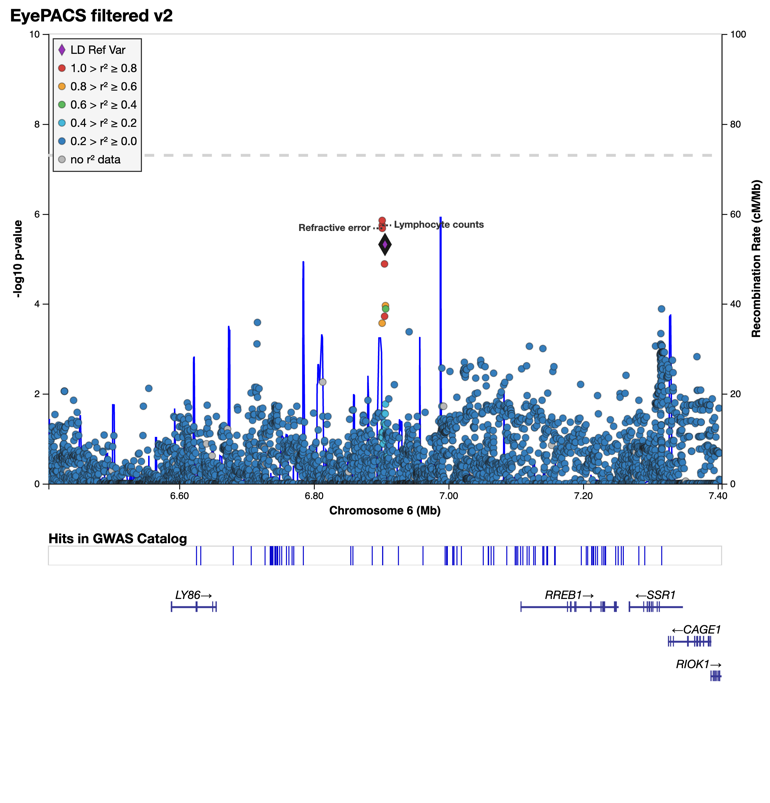 |
| Locus 21: chr6:151.047-151.547Mb - **ImageNet & EyePACS** (rs2064973/6:151297281_G/A) | 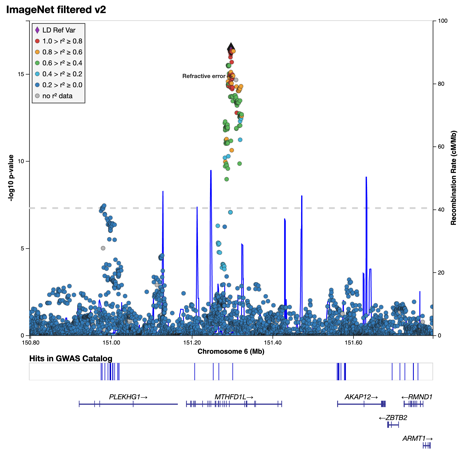 | 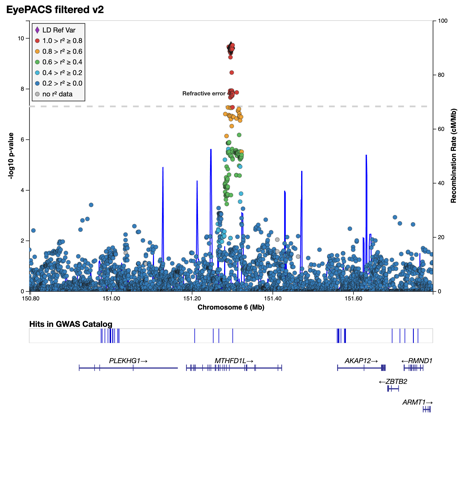 |
| Locus 22: chr7:96.216-96.716Mb - **ImageNet only** (rs13239805/7:96465530_T/C) | 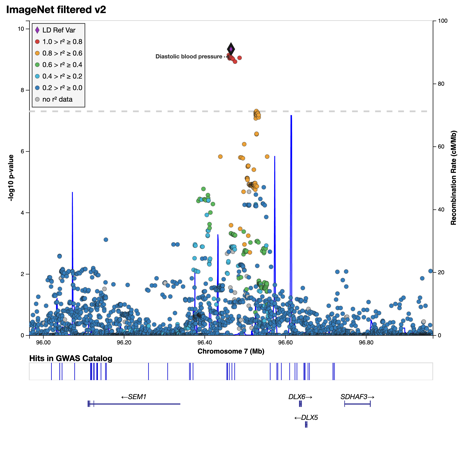 | 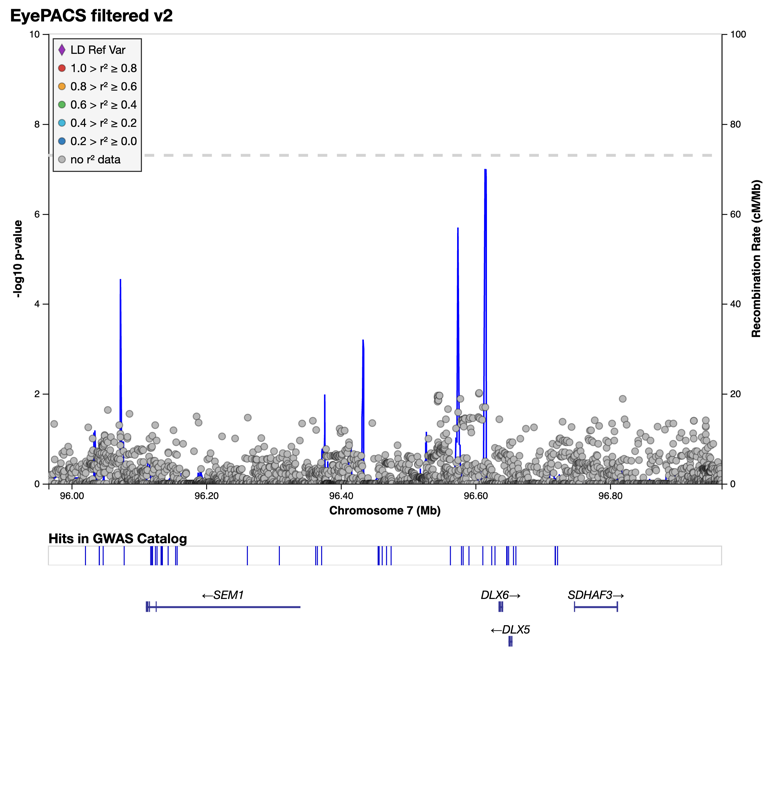 |
| Locus 23: chr7:98.717-99.696Mb - **ImageNet & EyePACS** (rs182206417/7:98967163_G/A) | 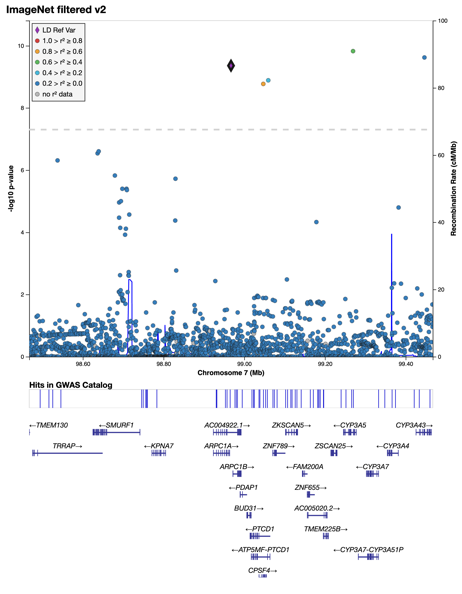 | 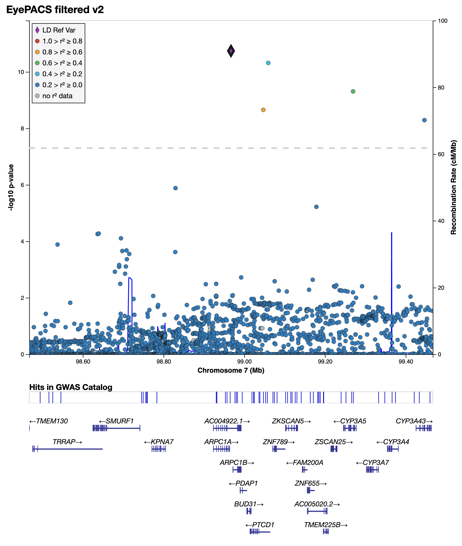 |
| Locus 23: chr7:98.717-99.696Mb - **ImageNet & EyePACS** (rs117445819/7:99269625_A/T) | 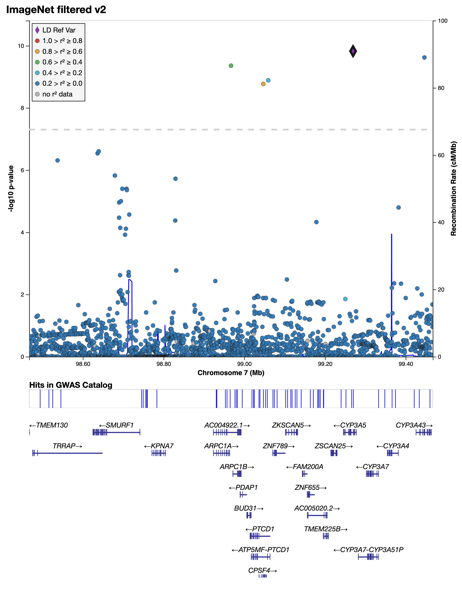 | 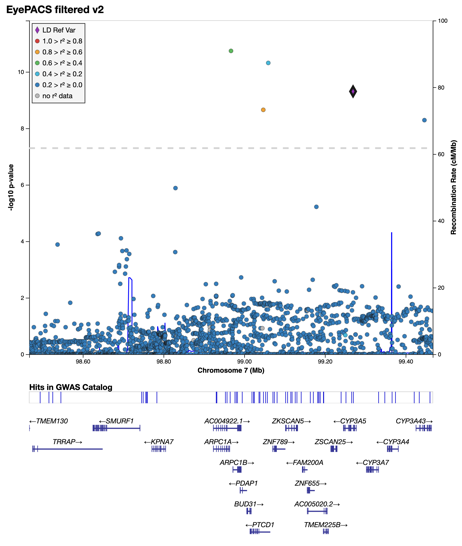 |
| Locus 23: chr7:98.717-99.696Mb - **ImageNet, EyePACS subsign** (rs117024920/7:99446494_C/A) | 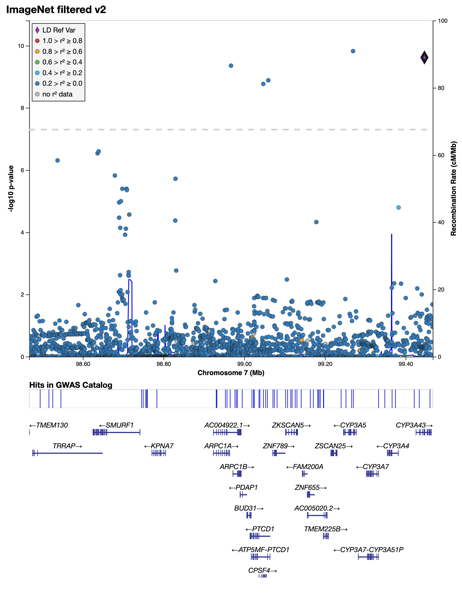 | 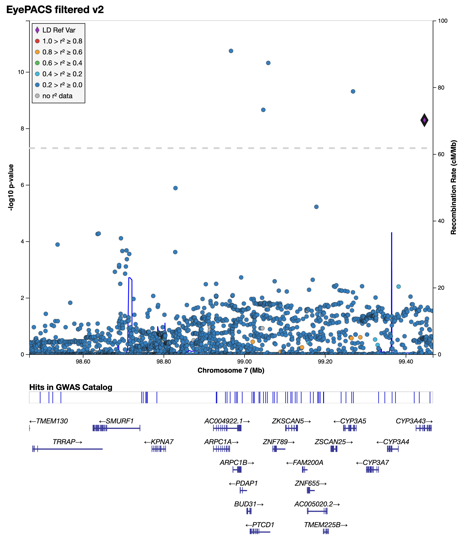 |
| Locus 24: chr7:100.141-100.913Mb - **ImageNet & EyePACS** (rs181872689/7:100390635_A/C) | 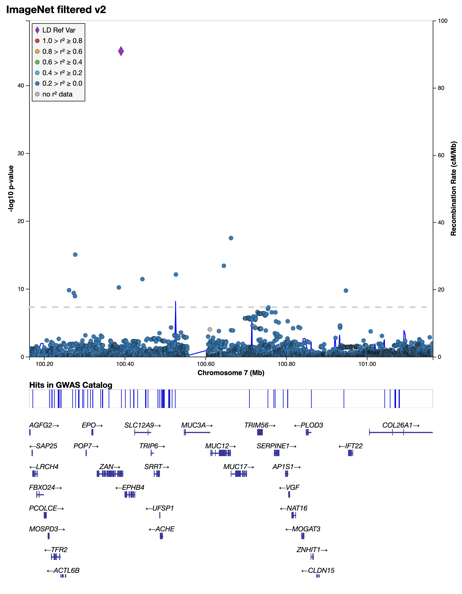 | 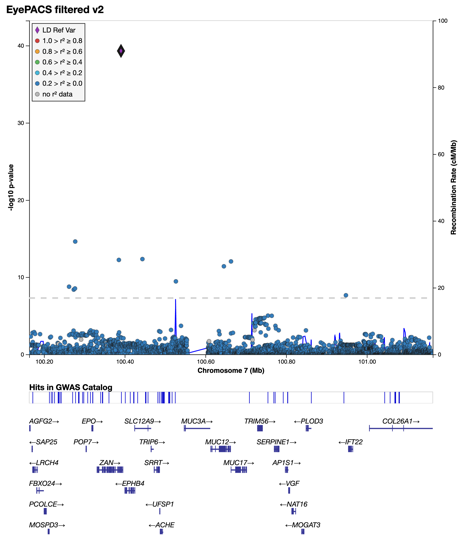 |
| Locus 24: chr7:100.141-100.913Mb - **ImageNet & EyePACS** (rs28690798/7:100663239_C/T) | 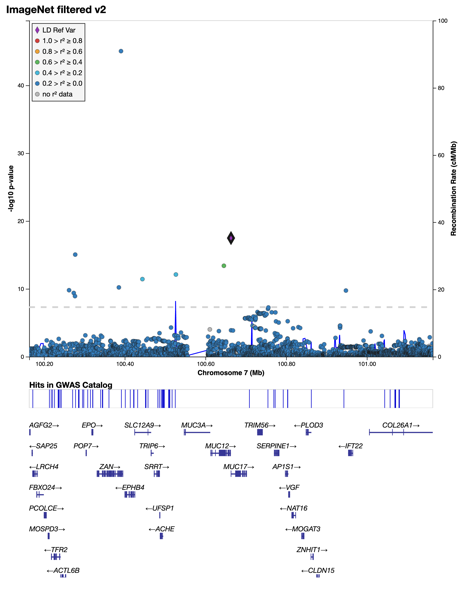 | 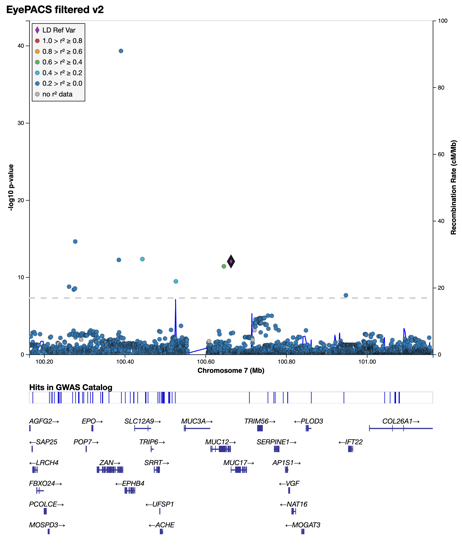 |
| Locus 25: chr7:100.914-101.414Mb  **EyePACS, ImageNet subsign** (rs149406562/7:101163592_C/A) | 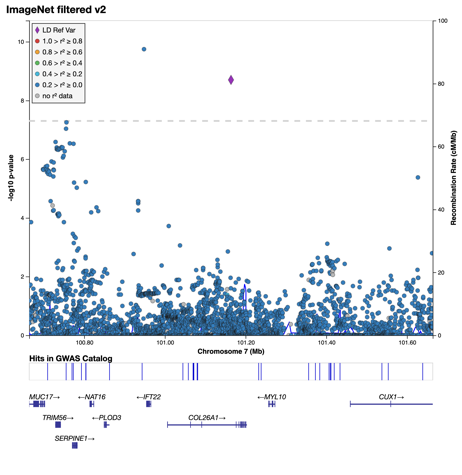 | 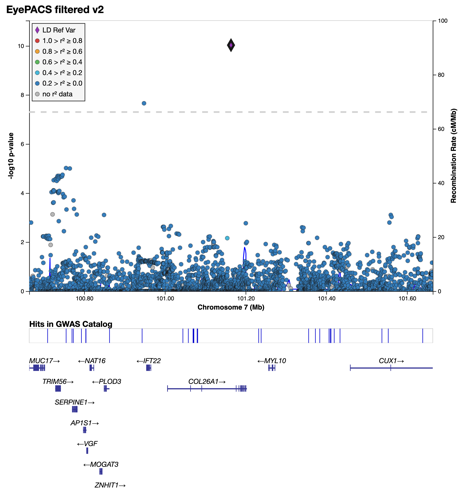 |
| Locus 26: chr8:95.774-96.274Mb  **EyePACS, ImageNet subsign**  (rs7832570/?) | 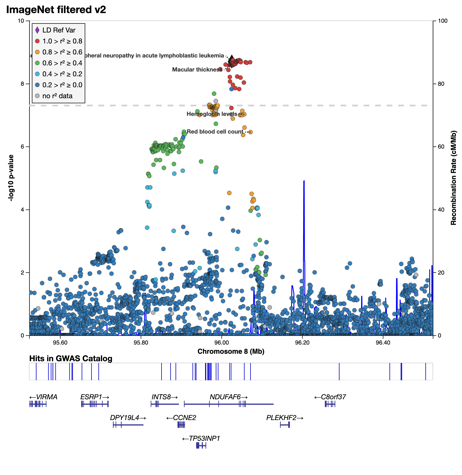 | 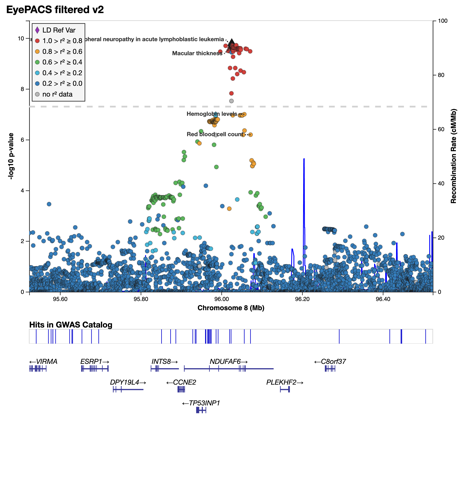 |
| Locus 27: chr8:108.877-109.715Mb - **ImageNet only** (rs34202872/8:109126735_C/CTTG) | 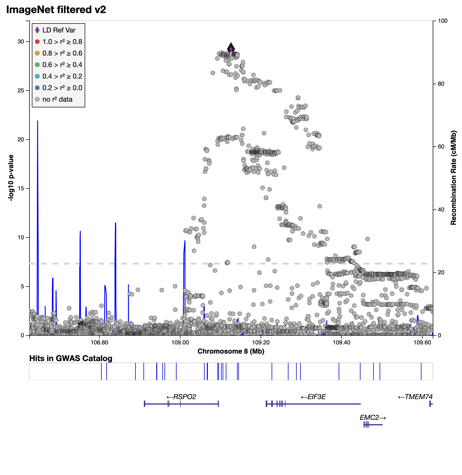 | 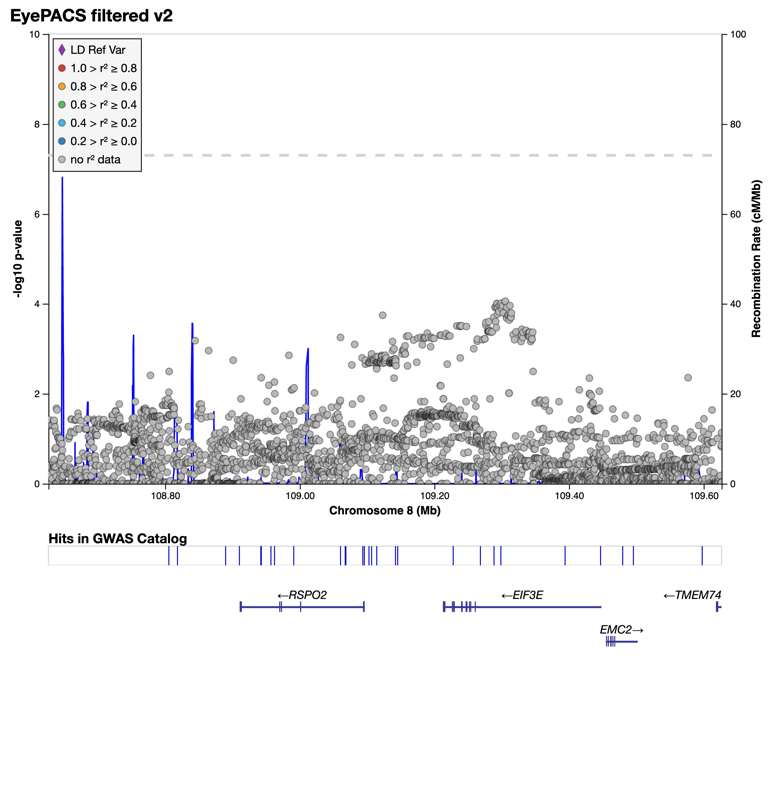 |
|  | 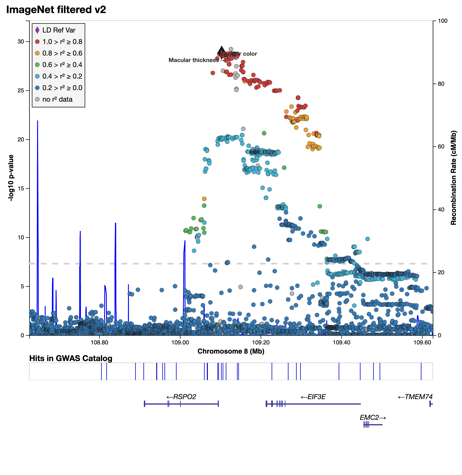  As there were no LD info for rs34202872 – used 8:109103640_A/G as well for LD ref |  |
| Locus 27: chr8:108.877-109.715Mb - **ImageNet only** (rs72668943/8:109465143_A/G) | 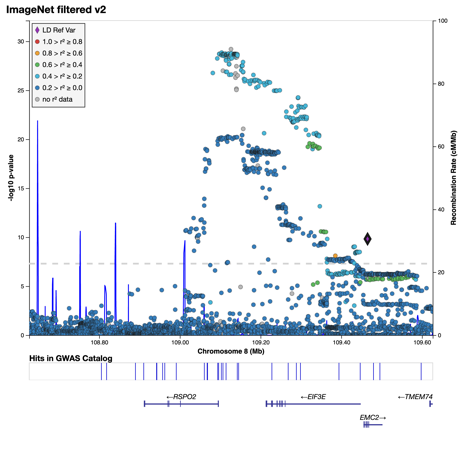 |  |
| Locus 28: chr9:12.425-12.925Mb  **ImageNet & EyePACS**  (rs10960752/9:12675284_G/A) | 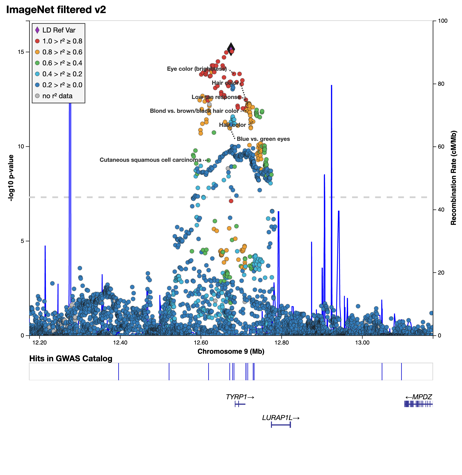 | 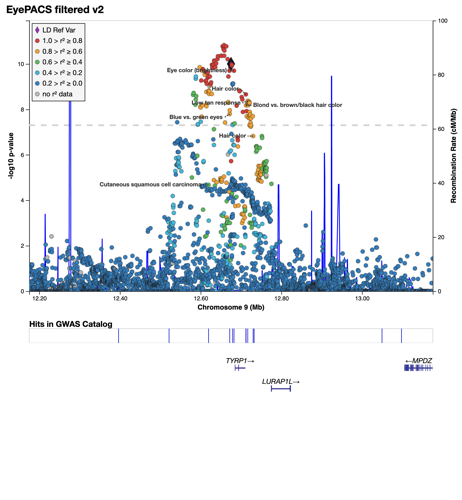 |
| Locus 29: chr9:130.086-130.586Mb - **ImageNet only** (rs35326378/9:130335728_C/CTTT) | 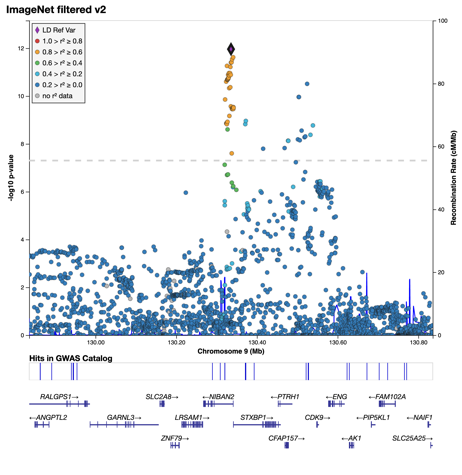 | 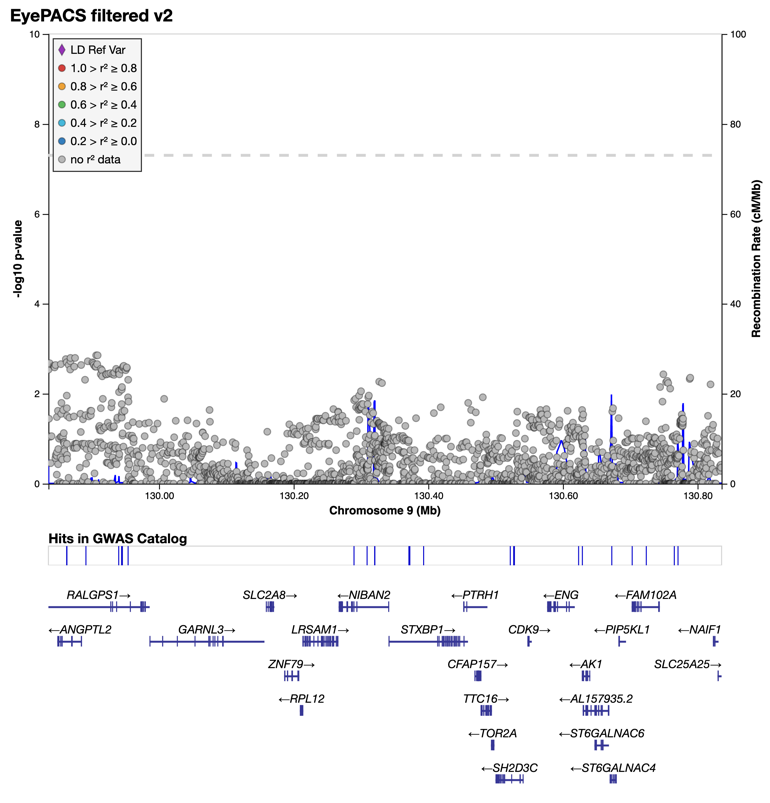 |
| Locus 30: chr10:76.968-77.468Mb  **ImageNet only**  (rs58628285/10:77218226_G/GCCCT) | 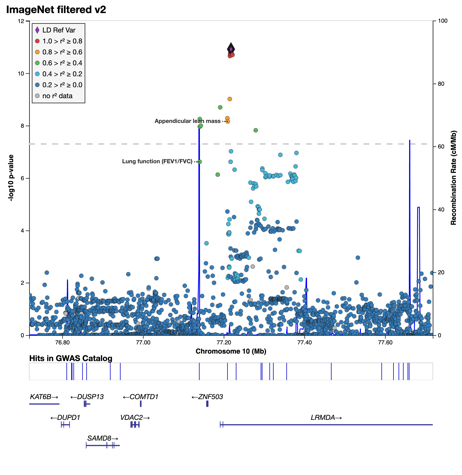 | 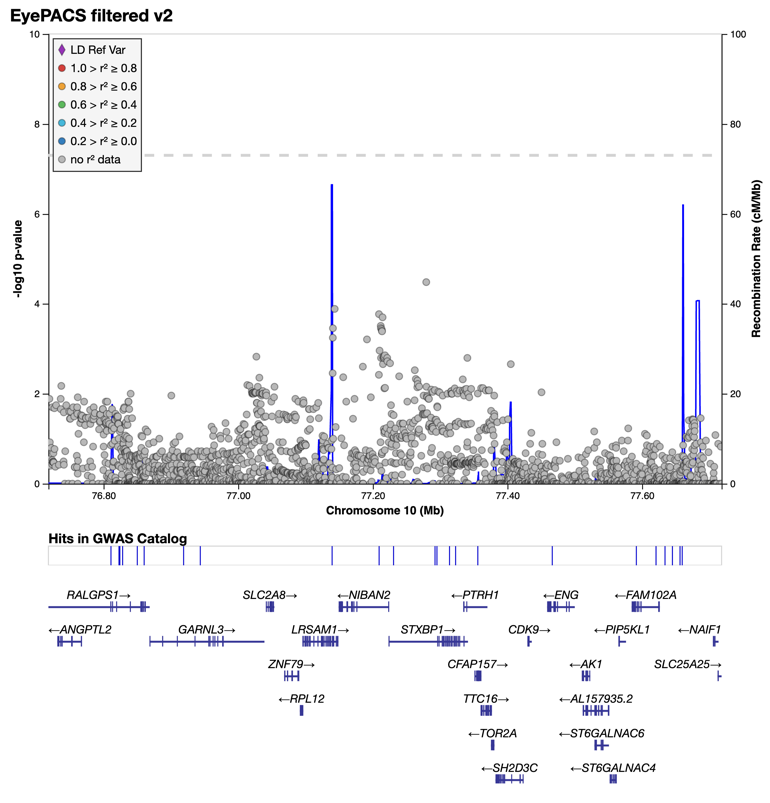 |
| Locus 31: chr11:15.976-16.476Mb  **ImageNet, EyePACS subsign** (rs7125634/11:16225741_C/T) | 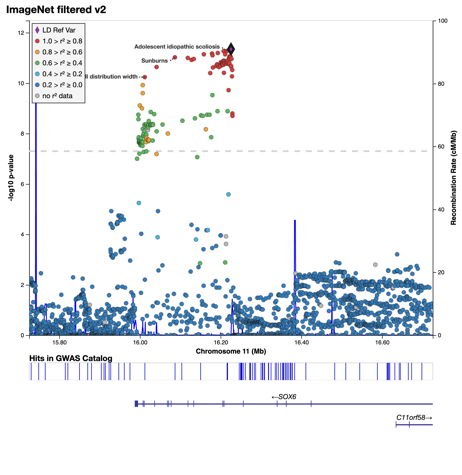 | 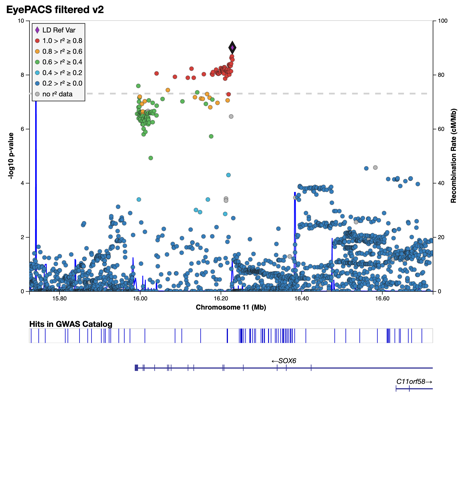  *11:16229139_CA/C used as LD ref |
| Locus 32: chr11:68.568-69.329Mb  **ImageNet & EyePACS**  (rs150527451/11:68817897_G/A) | 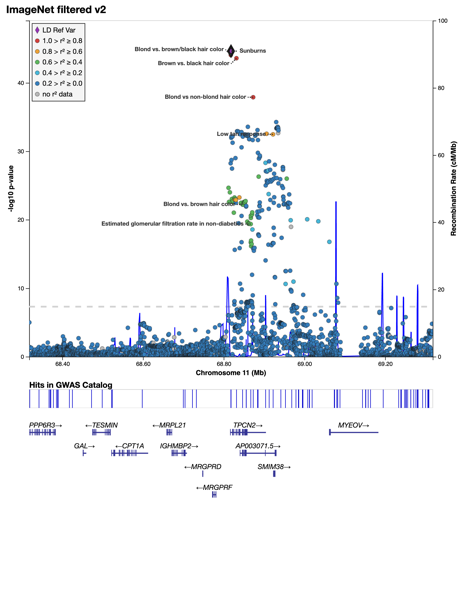 | 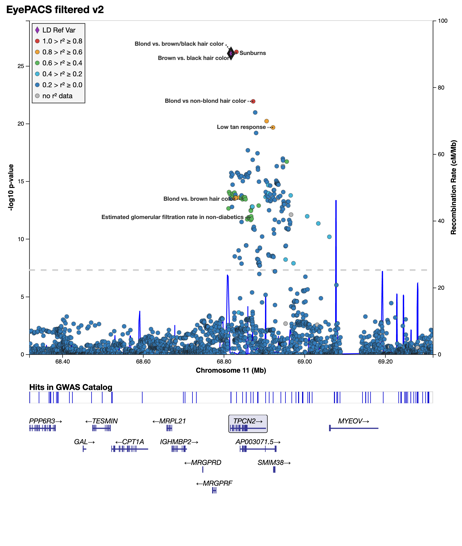 |
| Locus 32: chr11:68.568-69.329Mb  **ImageNet, EyePACS subsign**  (rs61746574/11:68840399_G/A) | 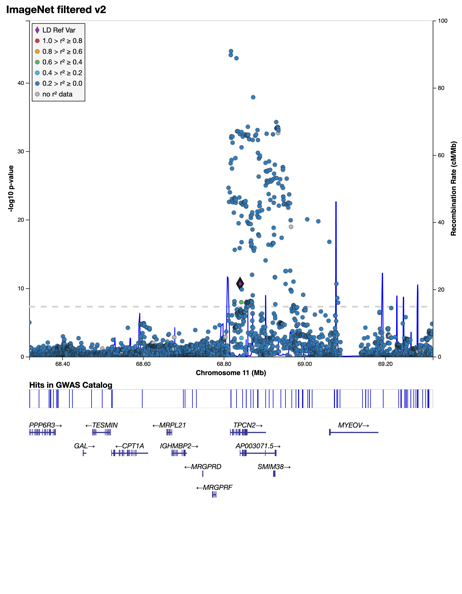 | 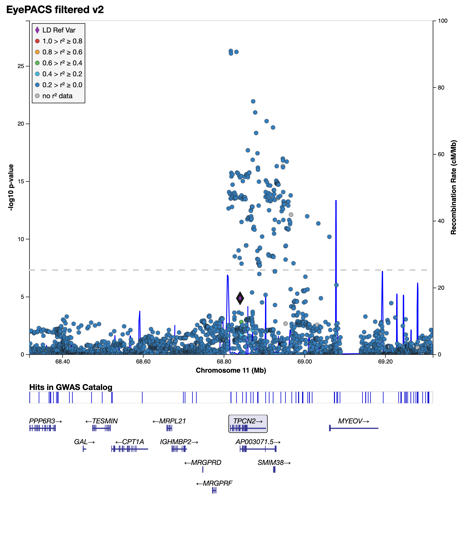 |
| Locus 32: chr11:68.568-69.329Mb  **ImageNet, EyePACS subsign**  (rs2924528/11:68926593_T/C) | 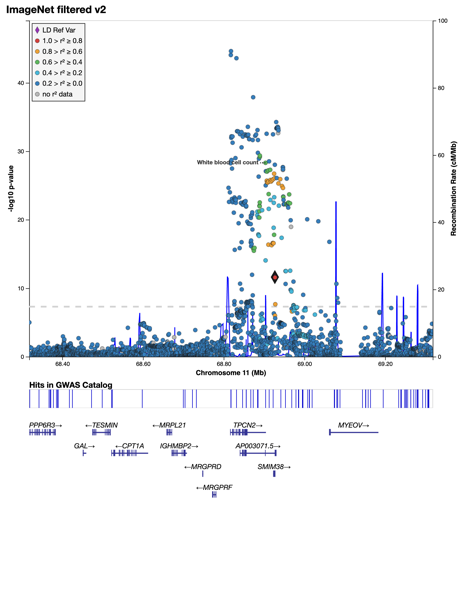 |  |
| Locus 32: chr11:68.568-69.329Mb  **ImageNet & EyePACS** (11:68929949_ATATTT_A/11:68929949_ATATTT/A) |  |  |
| Locus 32: chr11:68.568-69.329Mb - **ImageNet, EyePACS subsign** (rs12789955/11:69079134_T/G) |  |  |
| Locus 33: chr11:69.752-70.252Mb  **ImageNet & EyePACS** (rs751690958/11:70002184_TA/T) |  |  |
| Locus 34: chr11:87.644-88.144Mb - **ImageNet, EyePACS subsign** (rs188559/11:87893861_A/G) |  |  |
| Locus 35: chr11:88.366-89.268Mb - **ImageNet, EyePACS subsign** (rs151115969/11:88615937_G/A) |  |  |
| Locus 35: chr11:88.366-89.268Mb - **ImageNet, EyePACS subsign** (rs7942972/11:88708828_A/T) |  |  |
| Locus 35: chr11:88.366-89.268Mb  **ImageNet, EyePACS subsign** (rs3936623/11:88990049_T/C) |  | EyePACS P= 0.00077 |
| Locus 35: chr11:88.366-89.268Mb  **ImageNet, EyePACS subsign** (rs147546939/11:89011733_A/G) |  |  |
| Locus 35: chr11:88.366-89.268Mb  **ImageNet & EyePACS**  (rs1126809/11:89017961_G/A) |  |  |
| Locus 36: chr11:110.64-111.14Mb  **EyePACS, ImageNet subsign**  (rs317517/?) |  |  |
| Locus 37: chr12:6.705-7.205Mb  **ImageNet & EyePACS**  (rs5442/12:6954864_G/A) |  |  |
| Locus 38: chr12:20.339-20.839Mb  **ImageNet & EyePACS** (rs11045245/12:20589390_A/G) |  |  |
| Locus 39: chr12:23.73-24.23Mb - **ImageNet only** (rs9971729/12:23979791_A/C) |  |  |
| Locus 40: chr12:55.866-56.366Mb  **ImageNet & EyePACS**  (rs3138142/12:56115585_C/T) |  |  |
| Locus 41: chr12:95.882-96.404Mb - **ImageNet only** (rs17288108/12:96131895_A/G) |  |  |
| Locus 41: chr12:95.882-96.404Mb - **ImageNet only** (12:96154458_CT_C/12:96154458_CT/C) |  |  |
| Locus 42: chr12:111.635-113.133Mb  **EyePACS, ImageNet subsign** (rs3184504/12:111884608_T/C) |  |  |
| Locus 42: chr12:111.635-113.133Mb - **EyePACS only** (rs2013002/12:112200150_T/C) |  |  |
| Locus 42: chr12:111.635-113.133Mb - **EyePACS only** (rs17630235/12:112591686_G/A) |  |  |
| Locus 42: chr12:111.635-113.133Mb - **EyePACS only** (rs11066309/12:112883476_G/A) |  |  |
| Locus 43: chr13:28.917-29.42Mb - **ImageNet, EyePACS subsign** (rs9508078/13:29167474_A/G) |  |  |
| Locus 43: chr13:28.917-29.42Mb - **ImageNet, EyePACS subsign** (rs9506022/13:29169648_C/G) |  |  |
| Locus 44: chr13:94.949-95.449Mb - **ImageNet & EyePACS** (rs35321205/13:95199377_C/CT) |  |  |
| Locus 45: chr13:110.788-111.335Mb  **EyePACS, ImageNet subsign** (rs80281875/13:111038374_T/C) |  |  |
| Locus 45: chr13:110.788-111.335Mb  **ImageNet & EyePACS** (rs9559797/13:111085411_G/C) |  |  |
| Locus 46: chr14:60.763-61.263Mb  **ImageNet only** (rs2351174/14:61013237_G/T) |  |  |
| Locus 47: chr14:74.333-74.833Mb - **ImageNet, EyePACS subsign** (14:74583005_CAAAAAAT_C/14:74583005_CAAAAAAT/C) |  |  |
| Locus 48: chr14:105.657-106.157Mb - **ImageNet only** (rs56130943/14:105906522_A/C) |  |  |
| Locus 49: chr15:27.591-28.785Mb  **ImageNet & EyePACS**  **Entire Region** |  |  |
| Locus 50: chr15:28.800-29.589Mb  **ImageNet & EyePACS** (rs56166703/15:29033909_G/A) |  |  |
| Locus 50: chr15:28.800-29.589Mb - **ImageNet & EyePACS** (15:29318172_AAGAAATGGCCC_A/15:29318172_AAGAAATGGCCC/A) |  |  |
| Locus 50: chr15:28.800-29.589Mb - **EyePACS, ImageNet subsign** (rs35615406/15:29339457_G/A) | Pval Imagenet = 0.0001 |  |
| Locus 51: chr15:34.756-35.256Mb - **ImageNet & EyePACS** (rs634990/15:35006073_T/C) |  |  |
| Locus 52: chr15:74.82-75.32Mb - **ImageNet, EyePACS subsign** (rs12905199/15:75070196_A/G) |  |  |
| Locus 53: chr15:89.511-90.011Mb - **EyePACS, ImageNet subsign** (rs2070780/15:89760997_C/T) |  |  |
| rs2070780 / 15:89723378_G/A for comparison to rs2070780 |  |  |
| Locus 54: chr17:19.173-19.673Mb - **ImageNet, EyePACS subsign** (rs4924997/17:19422725_A/T) |  |  |
| Locus 55: chr17:68.095-68.595Mb - **ImageNet, EyePACS subsign** (rs11658102/17:68345127_T/C) |  |  |
| Locus 56: chr17:70.14-70.64Mb  **EyePACS only** (rs11077607/17:70390452_A/G) |  |  |
| Locus 57: chr17:79.365-79.865Mb  **ImageNet, EyePACS subsign**  (17:79614932_TTAAC_T/17:79614932_TTAAC/T) |  |  |
| Locus 58: chr19:38.9-39.4Mb  **ImageNet, EyePACS subsign**  (rs10415219/19:39150235_A/G) |  |  |
| Locus 59: chr19:48.979-49.479Mb - **EyePACS only** (rs479486/19:49229323_G/A) |  |  |
| Locus 60: chr20:57.555-58.055Mb - **ImageNet only** (rs118170335/20:57804695_C/T) |  |  |
